# KIBRA repairs synaptic plasticity and promotes resilience to tauopathy-related memory loss

**DOI:** 10.1101/2023.06.12.543777

**Authors:** Grant Kauwe, Kristeen A. Pareja-Navarro, Lei Yao, Jackson H. Chen, Ivy Wong, Rowan Saloner, Helen Cifuentes, Alissa L. Nana, Samah Shah, Yaqiao Li, David Le, Salvatore Spina, Lea T. Grinberg, William W. Seeley, Joel H. Kramer, Todd C. Sacktor, Birgit Schilling, Li Gan, Kaitlin B. Casaletto, Tara E. Tracy

## Abstract

Synaptic plasticity is obstructed by pathogenic tau in the brain, representing a key mechanism that underlies memory loss in Alzheimer’s disease (AD) and related tauopathies. Here, we define a mechanism for plasticity repair in vulnerable neurons using the C-terminus of the KIdney/BRAin (KIBRA) protein (CT-KIBRA). We show that CT-KIBRA restores plasticity and memory in transgenic mice expressing pathogenic human tau; however, CT-KIBRA did not alter tau levels or prevent tau-induced synapse loss. Instead, we find that CT-KIBRA binds to and stabilizes protein kinase Mζ (PKMζ) to maintain synaptic plasticity and memory despite tau mediated pathogenesis. In humans we find that reduced KIBRA in brain and increased KIBRA in cerebrospinal fluid are associated with cognitive impairment and pathological tau levels in disease. Thus, our results distinguish KIBRA both as a novel biomarker of synapse dysfunction in AD and as the foundation for a synapse repair mechanism to reverse cognitive impairment in tauopathy.

## INTRODUCTION

Pathological tau protein accumulates and forms aggregates in the brain in neurodegenerative diseases classified as tauopathies, including Alzheimer’s disease (AD), Pick’s disease, corticobasal degeneration (CBD) and progressive supranuclear palsy (PSP). Aberrant tau posttranslational modifications (PTMs) found in tauopathy brains can alter tau protein function, promote tau aggregation, and trigger toxicity in cells (1, 2). The extent of pathologically modified tau that accumulates in human brain correlates with dementia severity in disease (3, 4). Tau is predominantly localized within axons of healthy neurons, whereas an abundance of pathological tau with PTMs is found at synapses in AD brain (5–7). Synapse dysfunction is one of the earliest pathophysiological changes in tauopathy mouse models that precedes neurodegeneration and coincides with the start of cognitive impairments (8–10). Long-term potentiation (LTP) is an important plasticity mechanism at synapses in the hippocampus that underlies the formation of new memories, and LTP is inhibited by tau with mutations that cause familial frontotemporal dementia (8, 10, 11), as well as several AD-associated pathogenic forms of tau including hyperacetylated tau (9), hyperphosphorylated tau (12, 13), and tau oligomers (14, 15). Synapses, and their ability to express LTP, are particularly vulnerable to tau-induced toxicity in the brain.

KIdney/BRAin (KIBRA) is a postsynaptic protein encoded by the *WWC1* gene that has a single nucleotide polymorphism linked to memory and risk of late-onset AD in humans (16–19). The KIBRA protein is required for hippocampal LTP and memory in mice (20), and it contains multiple functional domains acting as a postsynaptic scaffold with approximately twenty binding partners identified (21, 22), supporting a critical role for KIBRA in modulating synaptic signaling and strength. In humans, KIBRA protein is significantly diminished in brain tissue of individuals with severe AD dementia and reduced KIBRA levels are associated with abnormal hyperacetylated tau (9). Moreover, a mimic of hyperacetylated human tau expressed in transgenic mice (tauKQ^high^ mice) obstructs LTP by reducing KIBRA levels at synapses (9), supporting that the loss of KIBRA function at synapses underlies AD-related plasticity and memory impairments.

Here, we quantified cerebrospinal (CSF) KIBRA levels for the first time in humans and show that elevated KIBRA in CSF and reduced KIBRA levels in human brain correlate with pathogenic tau levels and dementia severity in adults with tauopathy. To investigate how KIBRA-mediated signaling modulates synapses in neurons with pathogenic tau, we generated a truncated version of the KIBRA protein, comprised of the C-terminal domain (CT-KIBRA). We found that expression of CT-KIBRA in neurons of transgenic mice with pathogenic tau is sufficient to reverse synaptic plasticity and memory impairments associated with AD and related dementias. Together with the observation that tau-related pathology persists in the mouse brain, this provides evidence that CT-KIBRA promotes synaptic and cognitive resilience to tau toxicity. We further investigated the mechanistic effect of CT-KIBRA on LTP in neurons with pathogenic tau, finding that CT-KIBRA repairs synapse function through its interaction with protein kinase Mζ (PKMζ), an atypical protein kinase C (PKC) isoform that regulates postsynaptic AMPA-type glutamate receptors (AMPARs) and sustains the maintenance of LTP and memory (23–25). These results highlight the potential for a therapeutic strategy to re-establish KIBRA function in neurons, repair synapses, and reverse memory loss in tauopathy patients with altered KIBRA levels.

## RESULTS

### KIBRA levels in human brain and CSF are associated with pathological tau and cognitive impairment in older adults

To examine the relationship between KIBRA and pathogenic tau in different tauopathies, we performed immunoblot analyses on human brain homogenates from the middle temporal gyrus of control, AD and Pick’s disease cases characterized by neuropathology (Figure 1A and Supplemental Table 1). To assess the abnormal accumulation of acetylated tau in the brains, we used the MAb359 antibody which recognizes tau acetylated on K274 and labels pathological tau in most tauopathies, including both AD and Pick’s disease (9, 26). The elevated levels of K274 acetylation on soluble tau exhibited a strong relationship with lower KIBRA levels in the brain among the tauopathies cases (Figure 1B). Reduced KIBRA levels also correlated with increased soluble tau phosphorylation (p-tau181) assayed by the AT270 immunoreactivity relative to total soluble tau levels (Figure 1C), indicating that KIBRA downregulation is linked to both hyperacetylation and hyperphosphorylation of tau in the brain.

**Figure 1.**
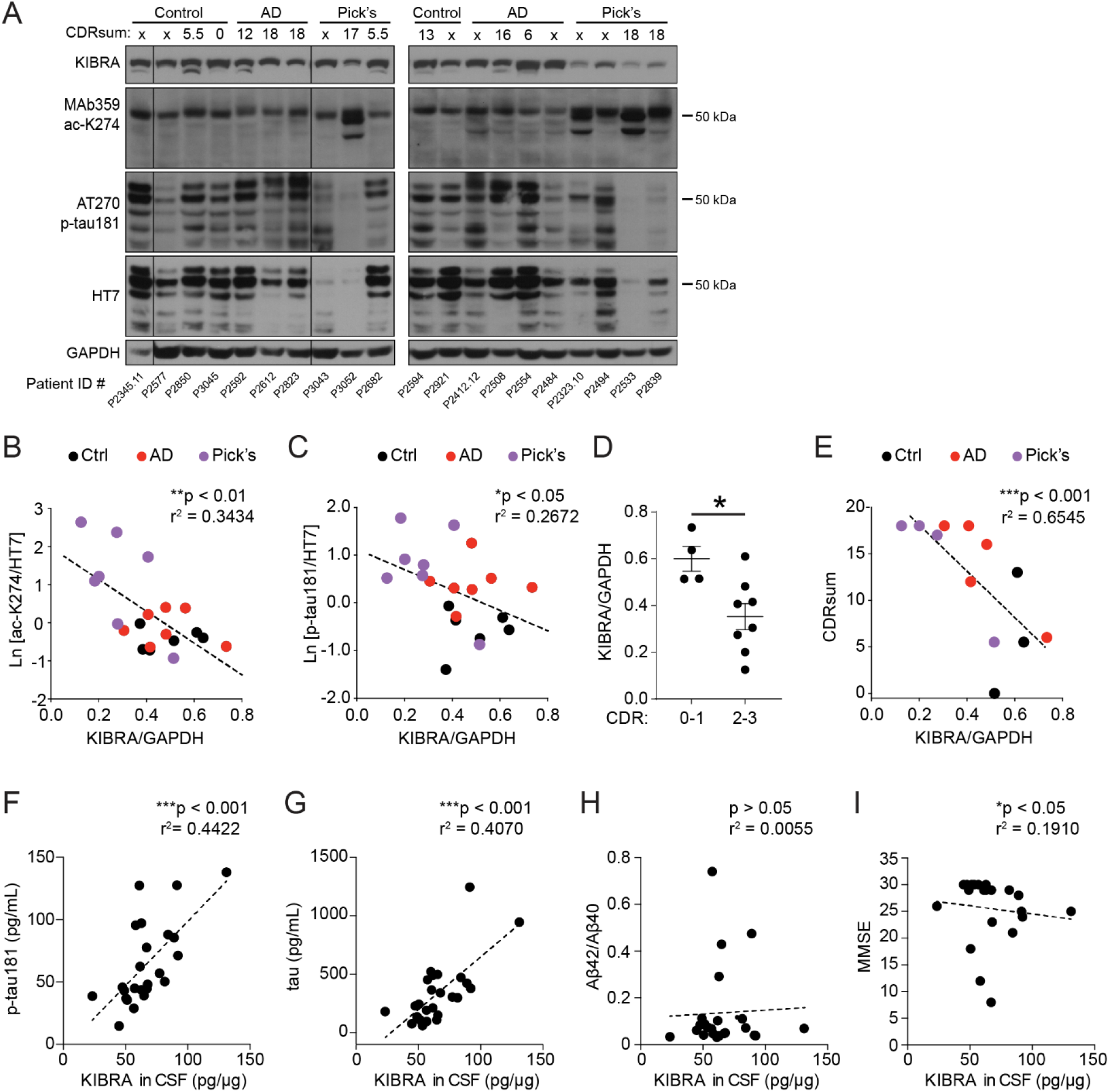
Reduced KIBRA levels in human brain and increased KIBRA levels in human CSF correlate with pathological tau levels and cognitive impairment in tauopathy. **(A)** Immunoblots of soluble homogenates from the middle temporal gyrus region of human brain classified as control, Alzheimer’s disease (AD) and Pick’s disease cases based on neuropathological diagnosis. Of the 20 cases analyzed, 12 were assigned a Clinical Dementia Rating (CDR) sum of boxes score based on cognitive function that are labeled above each lane. Lanes run on the same gel that were not adjacent are denoted by boundary lines. See also Supplemental Table 1. **(B and C)** Graph showing the relationship between KIBRA levels and the amount of soluble tau protein in human brains that is (**B**) acetylated on lysine 274 (ac-K274), recognized by the MAb359 antibody, and (**C**) phosphorylated on threonine 181 (p-tau181), recognized by the AT270 antibody (n = 6-7 cases/group; Spearman correlation). The immunoreactivity of acetylated and phosphorylated tau was quantified relative to total tau levels (HT7) which was measured from the respective immunoblots for MAb359 and AT270. **(D)** Graph of the KIBRA levels quantified in brain homogenate from the 12 cases that were assigned CDR scores (8 of the 20 brain samples used for this study were not assigned CDR scores). Cases with a CDR global score of 0-1, with none-to-mild dementia, were compared to CDR 2-3 cases, with moderate-to-severe dementia (n = 4-8 cases/group; * p < 0.05, unpaired Student’s *t*-test). Values are given as means ± SEM. **(E)** Analysis of the relationship between CDR sum of boxes score of neuropathological control, AD, and Pick’s disease cases and the levels of KIBRA in brain homogenate from 12 of the cases that were assigned CDR scores (n = 3-5 cases/group; Spearman correlation). **(F-H)** Spearman correlation analyses show the relationship between KIBRA levels and **(F)** p- tau181, **(G)** total tau, and **(H)** the Aβ42/Aβ40 ratio detected in individual CSF samples (n = 25 subjects for total tau analyses; n = 24 subjects for p-tau181 and Aβ42/Aβ40 ratio analyses). **(I)** Higher KIBRA levels in CSF are associated with worse cognition evaluated by Mini-Mental State Exam (MMSE, n = 21 subjects, Spearman correlation). See also Supplemental Figure 1 and Supplemental Table 2.

KIBRA levels are significantly diminished in the brain in adults with severe AD compared to non-demented controls (9). Here we explored the relationship between cognition and KIBRA in the brain across stages of dementia in tauopathy. Global Clinical Dementia Rating (CDR) scores were analyzed in twelve participants who ranged in dementia severity from none-to-mild (CDR 0-1) to moderate-to-severe dementia (CDR 2-3). Consistent with previous findings, KIBRA levels were significantly reduced in individuals with moderate-to-severe dementia (CDR 2-3) compared to those with none-to-mild (CDR 0-1) clinical impairment (Figure 1D). Comparisons were also made using the CDR sum of boxes (CDRsum) score, which allows for more continuous quantification of the extent of functional impairment compared to global CDR (27, 28). KIBRA levels strongly correlated with degree of functional severity (CDRsum) across control, AD and Pick’s disease brains, with the most impaired individuals having the lowest levels of KIBRA in the brain (Figure 1E).

Synaptic proteins can be detected in human CSF (29), including SNAP-25 (30) and neurogranin (31, 32), and are showing promise as biomarkers of *in vivo* synapse function in humans. Previous studies suggest that higher levels of several CSF synaptic proteins correlate with greater clinical dysfunction (33). To establish a method for detection of human KIBRA in CSF for the first time, we began by testing a KIBRA ELISA on lysates from HEK293T cells with or without KIBRA overexpression and we confirmed that the ELISA detected significant differences between KIBRA levels expressed at varying protein levels (Supplemental Figure 1A). We next used the ELISA to quantify the amount of KIBRA in CSF of 10 subjects with cognitive impairments due to AD, as well as 15 control subjects with normal cognition (Supplemental Table 2). KIBRA levels were compared to CSF tau protein including phosphorylated tau (p-tau181) and total tau which are both increased in CSF of AD patients and strongly correlate with worsened cognition (34). CSF KIBRA was significantly increased in subjects with clinically elevated CSF p-tau181 (Supplemental Figure 1B) (35). Furthermore, there was a strong correlation between higher KIBRA levels detected in CSF and increased p- tau181 and total tau levels (Figure 1, F and G). In contrast, KIBRA did not correlate with concentrations of Aβ40, Aβ42 or the ratio of Aβ42/Aβ40 in CSF (Figure 1H and Supplemental Figure 1, C and D), indicating that CSF KIBRA is predominantly associated with biomarkers of tau pathology rather than with Aβ pathology. To assess CSF KIBRA as a synaptic biomarker related to cognition in AD, we examined the relationship between CSF KIBRA levels and cognitive performances across participants on tasks of global cognition (Mini-Mental State Exam, MMSE). Higher CSF KIBRA levels significantly correlated with lower MMSE (Figure 1I). Together, these results demonstrate that reduced KIBRA in brain and increased KIBRA in CSF track strongly with clinical disease stage, cognitive impairment, and tau pathology both *in vivo* and postmortem in humans with tauopathy.

### A functional domain of the KIBRA protein reverses tauopathy-related synapse dysfunction

We next explored how KIBRA modulates the synaptic pathophysiology caused by pathogenic tau. To dissect the mechanistic effect of the KIBRA protein on tau-mediated synapse dysfunction, we generated constructs to express flag-tagged full-length human KIBRA or truncated forms of the KIBRA protein in neurons (Figure 2A). Two functional domains of the KIBRA protein were tested, one with 86 residues of the N-terminus containing the WW domains (NT-KIBRA) that interacts with synaptopodin and dendrin (36, 37), and the other with 187 residues of the KIBRA C-terminal domain (CT-KIBRA) that interacts with members of the PKC family including PKMζ (38–40). The constructs were expressed in cultured rat hippocampal neurons and immunostaining revealed that full-length human KIBRA and CT-KIBRA localized in dendritic spines, whereas the levels of NT-KIBRA in spines were significantly lower (Figure 2, B and C). We next examined the effect of the distinct KIBRA functional domains on the synaptic plasticity impairment caused by pathogenic tau. Neurons were transfected with either of the truncated KIBRA constructs together with human tau carrying lysine to glutamine mutations at lysine-274 and lysine-281 (tauKQ) to mimic hyperacetylated pathogenic tau that accumulates in the brain in AD and most other tauopathies (9, 26) . A chemical induction method, involving glycine treatment in the absence of magnesium, was applied to the neurons to induce LTP, which potentiates synapse function through recruitment of AMPARs to the surface of postsynaptic spines (41, 42). There was a significant increase in the surface immunolabeling of GluA1-containing AMPARs in spines in control neurons after chemical LTP induction whereas AMPAR insertion was obstructed in neurons expressing tauKQ with or without NT-KIBRA (Figure 2, D and E). Co-expression of CT-KIBRA with tauKQ in neurons rescued the increase in AMPARs on the spine surface triggered by chemical LTP (Figure 2, D and E). Likewise, the activity-induced recruitment of AMPARs to spines was blocked in neurons expressing human tau carrying the P301L mutation that causes familial frontotemporal dementia (FTD), and co expression of CT-KIBRA with the FTD mutant tau rescued AMPAR recruitment during chemical LTP (Supplemental Figure 2, A and B).

**Figure 2.**
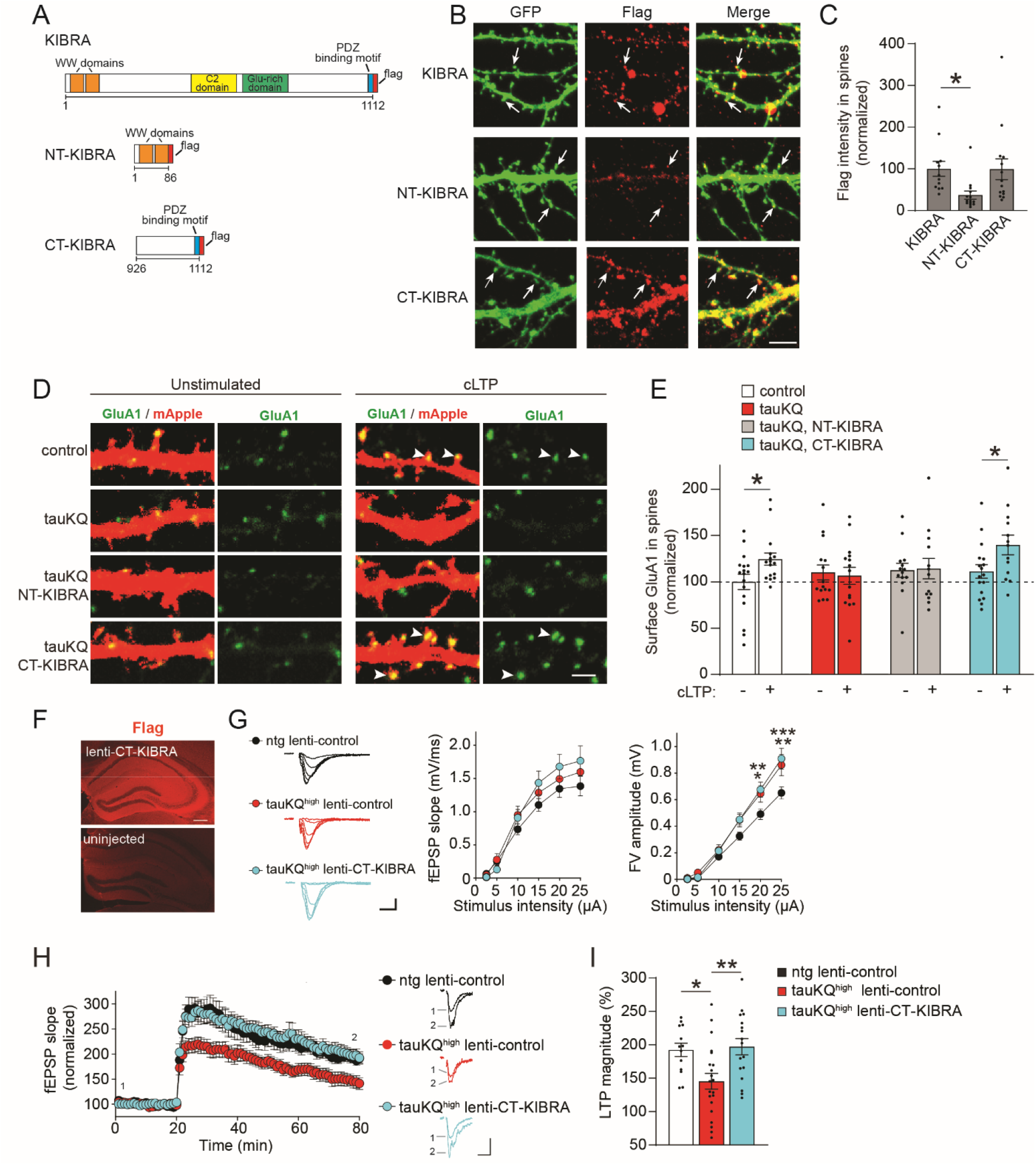
CT-KIBRA reverses the impairment of activity-dependent postsynaptic AMPAR trafficking and LTP caused by pathogenic tau. **(A)** Design of full-length KIBRA and truncated KIBRA constructs fused to a flag tag. NT-KIBRA contained the first 86 residues and CT-KIBRA included the last 187 residues of KIBRA. **(B)** Each flag-tagged KIBRA construct (red) was co-expressed with GFP (green) in cultured rat hippocampal neurons. Immunolabeling of neurons expressing each construct detected KIBRA, NT-KIBRA and CT-KIBRA in dendritic spines filled with GFP (arrows). Scale bar, 5 µm. **(C)** Quantification of the intensity of flag immunoreactivity in dendritic spines in neurons expressing full-length KIBRA, NT-KIBRA, or CT-KIBRA (n = 12-14 neurons/group; *p < 0.05, one-way ANOVA, Bonferroni *post*-hoc analyses). **(D)** Images of surface AMPAR immunolabeling (green), labeled with an N-terminal GluA1 antibody, in neurons expressing mApple (red) with or without tauKQ and KIBRA constructs. Neurons were either unstimulated or treated with a chemical LTP (cLTP) method to induce synaptic plasticity and AMPAR trafficking (arrowheads). Scale bar, 2 µm. **(E)** The mean intensity of surface GluA1 immunofluorescence in spines was quantified for unstimulated and cLTP-induced neurons (n = 13-17 neurons/group; *p < 0.05, unpaired Student’s *t*-test). See also Supplemental Figure 2. **(F)** Representative images of flag immunolabeling (red) detected in the ipsilateral hippocampus of a mouse injected with flag-tagged CT-KIBRA lentivirus with the control hippocampus on the contralateral side that was not injected with lentivirus. Scale bar, 200 µm. **(G)** TauKQ^high^ and ntg mice were injected with control empty vector lentivirus or CT-KIBRA lentivirus at 15-16 months old. Field recordings were performed in the dentate gyrus molecular layer of brain slices from mice at 17-18 months of age. The field excitatory postsynaptic potentials (fEPSP) slopes and the fiber volley (FV) amplitudes were measured in response to increasing stimulus intensities applied to the perforant pathway inputs. Representative traces and graphs of mean fEPSP slope and FV amplitude were generated from field recordings of slices from each group (n = 12-14 slices from four mice/group). TauKQ^high^ mouse brain slices exhibited significantly increased mean FV amplitude in dentate gyrus at the highest stimulus intensities (20 µA stimulus: ntg lenti-control vs. taukQ^high^ lenti-control *p < 0.05, ntg lenti-control vs. taukQ^high^ lenti-CT-KIBRA **p < 0.01; 25 µA stimulus: ntg lenti-control vs. taukQ^high^ lenti-control **p < 0.01, ntg lenti-control vs. taukQ^high^ lenti-CT-KIBRA ***p < 0.001, repeated measures two-way ANOVA, Bonferroni *post*-hoc analyses). Scale bars, 0.5 mV and 5 ms. **(H)** Representative traces of fEPSPs recorded in the dentate gyrus before and after TBS of the perforant pathway to induce LTP in dentate granule cells. Quantification of the fEPSP slopes 20 min before and 60 min after TBS was normalized to the average fEPSP slope during the first 20 min of the baseline recording for each slice (n = 13-21 slices from five to seven mice/group). Scale bars, 0.4 mV and 10 ms. **(I)** Graph of the mean fEPSP slopes at 55-60 min after TBS induction representing the magnitude of LTP (n = 13-21 slices from five to seven mice/group; *p < 0.05, **p < 0.01, one way ANOVA, Bonferroni *post*-hoc analyses). Values are given as means ± SEM.

Based on our finding that CT-KIBRA, but not NT-KIBRA, was sufficient to modulate AMPARs at synapses during plasticity, we next examined the effect of CT-KIBRA on LTP in aged transgenic mice expressing high levels of tauKQ (tauKQ^high^). TauKQ^high^ mice have impaired LTP in the dentate gyrus by 6 months of age (9). We generated lentivirus for flag tagged CT-KIBRA expression in mouse hippocampus following stereotaxic injection (Figure 2F). To determine the efficacy of CT-KIBRA in reversing synapse dysfunction in aged mice, the CT-KIBRA lentivirus (lenti-CT-KIBRA) or a control lentivirus (lenti-control) was bilaterally injected into the hippocampus of 15- to 16-month-old tauKQ^high^ mice and non-transgenic (ntg) controls. At 17 to 18 months of age, field recordings were performed in the dentate gyrus molecular layer with stimulation of the perforant pathway. Compared to ntg lenti-control mice, dentate granule cells in tauKQ^high^ mice with either lenti-control or lenti-CT-KIBRA exhibited similar basal postsynaptic responses to stimuli, but the fiber volley amplitudes recorded from perforant pathway projections into the dentate gyrus were significantly elevated at the highest stimulus intensities in both control and CT-KIBRA lentivirus treated tauKQ^high^ mice (Figure 2G). This suggests that tauKQ alters the maximum strength of perforant pathway inputs to dentate gyrus in aged mice, however this presynaptic effect of tauKQ was not modulated by CT-KIBRA. Induction of LTP by theta burst stimulation (TBS) of the perforant pathway triggered a persistent enhancement of the postsynaptic response in ntg lenti-control mice which was impaired in tauKQ^high^ lenti-control mice (Figure 2, H and I). The magnitude of LTP in dentate granule cells was significantly increased in tauKQ^high^ lenti-CT-KIBRA compared to tauKQ^high^ lenti-control mice, and CT-KIBRA enhanced the persistent strengthening of synapses in tauKQ^high^ mice to the same level as ntg lenti-control mice (Figure 2I). These results support that the C-terminal domain of the KIBRA protein is sufficient to restore AMPAR trafficking and LTP at synapses in hippocampal neurons with pathogenic tau.

### CT-KIBRA improves hippocampus-dependent memory in mice with pathogenic tau despite tauopathy-related pathology

We next assessed the functional impact of CT-KIBRA on hippocampus-dependent memory impairment caused by pathogenic tau in aged mice. TauKQ^high^ mice have pattern separation memory loss by 6-7 months of age coinciding with the LTP deficit in dentate gyrus (9). To test the impact of CT-KIBRA treatment after the onset of memory impairment in aged tauKQ^high^ mice, injections of lenti-CT-KIBRA or lenti-control into bilateral hippocampi were performed on tauKQ^high^ mice and ntg littermates at 10-12 months old and behavior tests were performed 4-6 weeks later. During the sample phase of the object-context discrimination test of pattern separation memory, the mice spent equivalent time exploring the distinct pairs of identical objects in two different but similar contexts (Figure 3A). In the memory test phase, one of the objects in each pair is switched between the contexts, thus presenting the mice with a familiar object that is incongruent with the context. The ntg mice explored the incongruent object significantly longer than the congruent object (Figure 3B), which is an indication of hippocampus-dependent pattern separation memory. The tauKQ^high^ lenti-control mice demonstrated impaired pattern separation memory, however lenti-CT-KIBRA restored the performance of tauKQ^high^ mice in discrimination of the mismatched object (Figure 3B).

**Figure 3.**
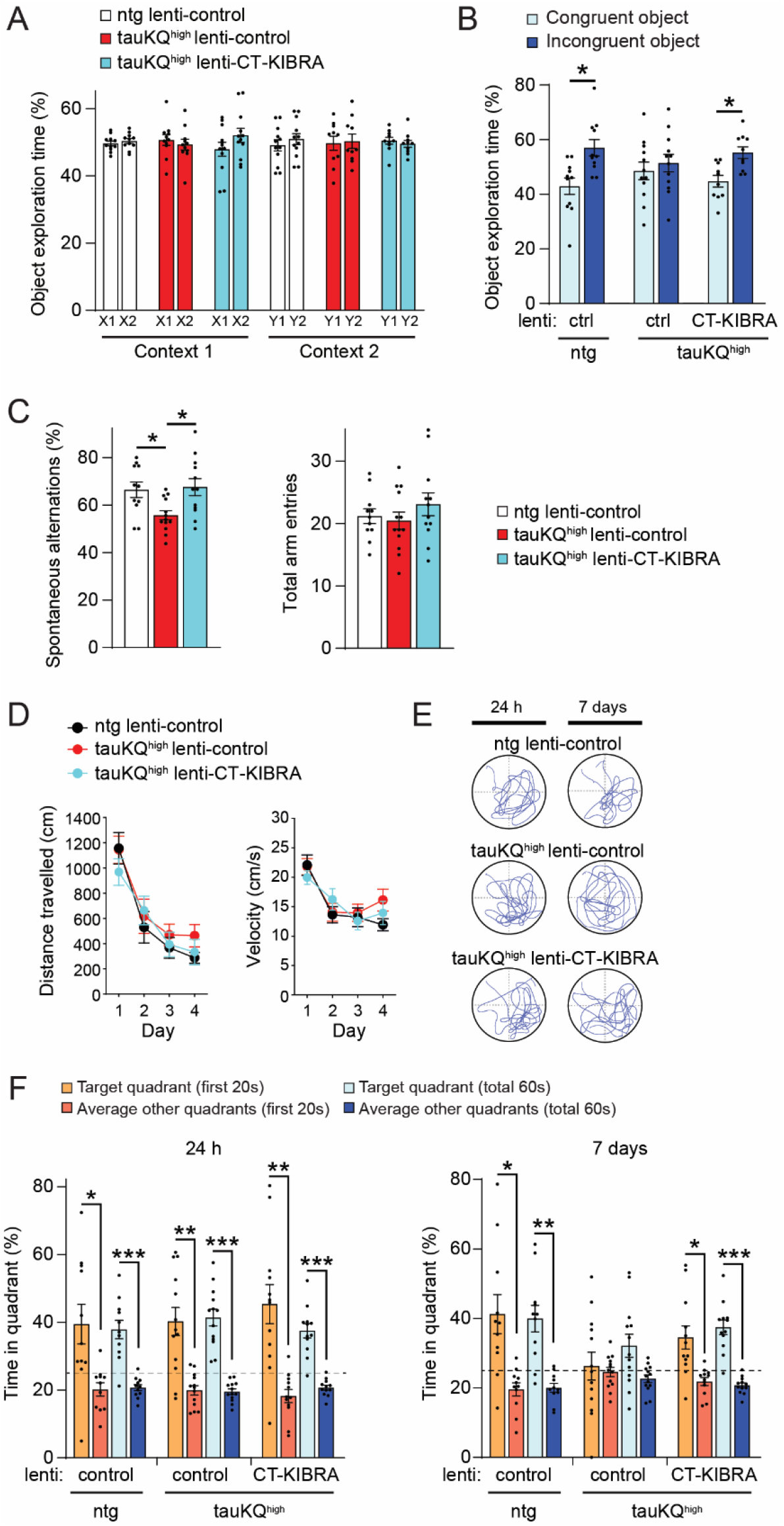
CT-KIBRA expression in hippocampus reverses memory impairment in tauKQ^high^ mice. **(A and B)** The object-context discrimination test was used to assess hippocampus-dependent pattern separation memory in mice (n = 10-12 mice/group). **(A)** During the sample phase, the mean proportion of time that the mice spent exploring two identical objects in each context was calculated. Each mouse explored the identical X1 and X2 objects in Context 1, and the identical Y1 and Y2 objects in Context 2 which were distinct from the objects in Context 1. **(B)** The mean percent time spent exploring the incongruent and congruent objects in both contexts for each group was analyzed during the test phase of the object-context discrimination test (*p < 0.05, paired Student’s *t*-test). **(C)** Hippocampus-dependent spatial working memory was monitored from ntg and tauKQ^high^ mice in the Y-maze test (n = 11-13 mice/group). Graph of the mean percentage of spontaneous alternations completed by the mice in the Y-maze within 5 min (left, *p < 0.05, one-way ANOVA, Bonferonni *post-*hoc analyses). The average number of total Y-maze arm entries made by the mice (right) was equivalent across each group. **(D-G)** Mice were tested for spatial learning and memory in the Morris water maze (n = 11-13 mice/group). **(D)** The mean distance travelled to the hidden platform and the average swim velocity were measured during hidden platform training (p > 0.05, repeated measures two-way ANOVA). **(E)** Representative swim paths of mice during the probe trials for spatial memory testing performed at 24 h and 7 days after hidden platform training. (**F**) Graphs of the mean percent time spent in the target quadrant compared to the average time spent in the other quadrants in the 24 hr (left) and 7-day (right) probe trials. Quantification and comparisons between target quadrant and other quadrants were analyzed for both the first 20 s of the probe trial and for the total 60 s period of the probe trial for all mice (*p < 0.05, **p < 0.01, and ***p < 0.001, paired Student’s *t*-test). In the 7-day probe trial, tauKQ^high^ lenti-control mice did not spend significantly more time in the target quadrant either in the first 20 s or in the total 60 s period of the probe trial. The dotted line marks the 25% chance level. Values are given as means ± SEM.

The tauKQ^high^ mice were next tested in the Y-maze to determine whether CT-KIBRA has an impact on hippocampus-dependent working memory. TauKQ^high^ lenti-control mice demonstrated impaired working memory, whereas the proportion of spontaneous alternations completed by tauKQ^high^ lenti-CT-KIBRA mice was significantly increased without affecting the total number of arm entries during the test (Figure 3C). Thus, CT-KIBRA improved spatial working memory in tauKQ^high^ mice. In the Morris water maze (MWM) test of spatial learning and memory, ntg lenti control, taukQ^high^ lenti-control, and tauKQ^high^ lenti-CT-KIBRA mice showed comparable swim speeds and spatial learning to find the hidden platform during training (Figure 3D). The ntg lenti control, taukQ^high^ lenti-control, and tauKQ^high^ lenti-CT-KIBRA mice spent a greater fraction of time in the target quadrant in the probe trial 24 h after the hidden platform training was completed (Figure 3, E and F). Only the ntg lenti-control and tauKQ^high^ lenti-CT-KIBRA mice maintained a significant preference for the target quadrant probe trial after 7 days, whereas tauKQ^high^ lenti-control mice did not (Figure 3, E and F), indicating that CT-KIBRA reversed the long-term spatial memory impairment in tauKQ^high^ mice. Overall, these results support that CT-KIBRA enhances multiple forms of hippocampus-dependent memory that are compromised by pathogenic tau in the brain.

We next examined whether CT-KIBRA mitigates pathology in the brain caused by expression of the acetylated tau mimic. CT-KIBRA did not prevent the accumulation of tau phosphorylated at serine 202/205 (AT8) that was significantly increased in mossy fibers in the hippocampus of tauKQ^high^ mice (Figure 4, A-C). Total levels of human tau and tau phosphorylation at threonine 231 (AT180) and threonine 181 (AT270) in the hippocampus of tauKQ^high^ mice were also not altered by CT-KIBRA expression (Figure 4, D and E). Synaptic vesicle associated protein 2 (SV2) has been used as a marker of synapse loss in human AD brain (43, 44). To assess synapse deterioration in tauKQ^high^ mice, we analyzed immunolabeling of SV2 and found that it was significantly decreased in the CA1 region of the hippocampus in tauKQ^high^ with or without CT-KIBRA compared to ntg controls (Figure 4, F and G). Thus, while CT-KIBRA was sufficient to restore hippocampal synaptic plasticity, it was not sufficient to reverse tau-mediated synapse loss in the CA1 region. SV2 immunolabeling in the molecular layer of the dentate gyrus was not altered in tauKQ-expressing mice compared to controls (Supplemental Figure 3, A and B). Mass spectrometry analysis of protein extracted from whole hippocampus of tauKQ^high^ lenti-control and tauKQ^high^ lenti-CT-KIBRA mice was performed to monitor changes in the global proteome compared to ntg lenti-control mice (Supplemental Figure 3C and Supplemental Table 3). Notably, of the 64 hits identified that were either significantly increased or decreased in tauKQ^high^ compared to ntg mice, 33% of them changed in the hippocampi of both tauKQ^high^ lenti-control and tauKQ^high^ lenti-CT-KIBRA mice. A comparison of tauKQ^high^ mice with or without CT-KIBRA also yielded no significant differences (Q < 0.05) in the abundance of the proteins that were identified (Supplemental Table 3). Cellular component classification of proteins that were significantly downregulated in both tauKQ^high^ lenti-control or tauKQ^high^ lenti-CT-KIBRA mice compared to ntg lenti-control mice revealed similar gene ontology terms including synapses and postsynaptic sites (Supplemental Figure 3, D and E, and Supplemental Table 3). Proteins that were upregulated in tauKQ^high^ lenti-control and tauKQ^high^ lenti-CT-KIBRA mice compared to ntg lenti-control mice included microtubule and cytoskeleton components as well as axonal proteins (Supplemental Figure 3F and Supplemental Table 3). These findings suggest that CT-KIBRA restores synaptic plasticity and reverses tau-induced memory impairment without altering levels of pathological tau in the brain and without recovering CA1 synapses lost in the hippocampus.

**Figure 4.**
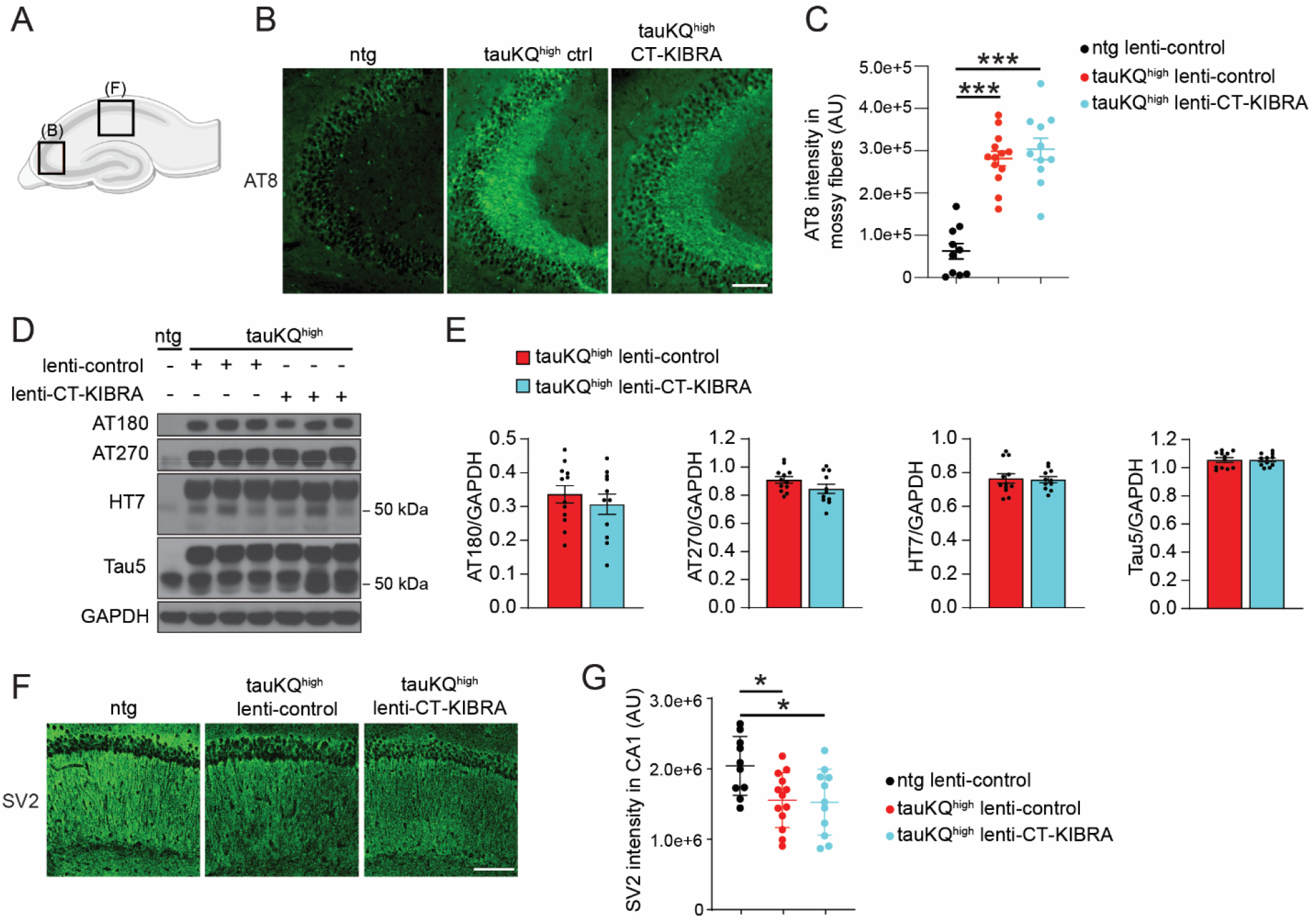
Elevated pathological tau and CA1 synapse loss persist in hippocampus of tauKQ^high^ mice with CT-KIBRA. **(A)** Illustration of a mouse hippocampus depicting where the imaging of mossy fibers in CA3 **(B)**nd the imaging of synapses in CA1 **(F)** was performed. **(B)** Representative images of AT8 immunostaining demonstrating accumulation of phosphorylated tau in the mossy fibers inputs to CA3 in tauKQ^high^ mice with or without CT KIBRA. Scale bar, 100 µm. **(C)** Graph of the mean integrated intensity of AT8 immunofluorescence in the mossy fibers of CA3 (n = 10-13 mice/group, *** p < 0.001, one-way ANOVA, Bonferonni *post*-hoc analyses). **(D)** Representative immunoblots of phosphorylated tau (AT180 and AT270 antibodies), human tau (HT7 antibody), and total tau (Tau5 antibody) from hippocampal homogenates of ntg lenti control, tauKQ^high^ lenti-control and tauKQ^high^ lenti-CT-KIBRA mice. **(E)** Quantification of phosphorylated tau and total tau in hippocampal homogenates normalized to GAPDH (n = 10-13 mice/group, p > 0.05, unpaired Student’s *t*-test). **(F)** Representative confocal images of SV2 immunolabeling as a marker of synapses in CA1. Scale bar, 100 µm. **(G)** Quantification of the mean integrated intensity of SV2 immunofluorescence in the stratum radiatum of CA1 (n = 10-13 mice/group; * p < 0.01, one-way ANOVA, Bonferonni *post*-hoc analyses). Values are given as means ± SEM. See also Supplemental Figure 3 and Supplemental Table 3.

### CT-KIBRA stabilizes and enhances PKM**ζ** at synapses with pathogenic tau

Previous studies established that KIBRA binds to PKMζ and prevents PKMζ degradation (38, 45). KIBRA also interacts with PICK1, which modulates AMPAR trafficking during plasticity (20). To examine the interaction of CT-KIBRA with PKMζ and PICK1, we used a proximity ligation assay (PLA) which detects protein-protein interactors within close proximity (<40 nm) (46). When expressed in HEK293T cells, HA-tagged PKMζ exhibited significantly stronger PLA signal with both full-length KIBRA and CT-KIBRA compared to NT-KIBRA (Figure 5, A and B). These results are consistent with the requirement of the KIBRA C-terminus for PKMζ binding (38), and confirm that the binding affinities of PKMζ to CT-KIBRA and full-length KIBRA are comparable. The PLA signal intensity in HEK293T cells expressing HA-tagged PICK1 together with full length KIBRA was significantly higher than in cells co-expressing NT-KIBRA or CT-KIBRA (Supplemental Figure 4, A and B), demonstrating that CT-KIBRA has a weaker association with PICK1 than full-length KIBRA. Given the strong interaction between CT-KIBRA and PKMζ, we next tested the effect of CT-KIBRA on PKMζ degradation kinetics in HEK293T cells. When HEK293T cells expressing PKMζ alone were treated with cycloheximide (CHX) to block protein synthesis, PKMζ levels quickly declined over 48 h (Figure 5, C and E). Consistent with the stabilizing effect of KIBRA on PKMζ, CT-KIBRA co-expression increased the total amount of PKMζ in HEK293T cells, and degradation of PKMζ was significantly slower following CHX treatment in cells with CT-KIBRA compared to those that only expressed PKMζ (Figure 5, C-E).

**Figure 5.**
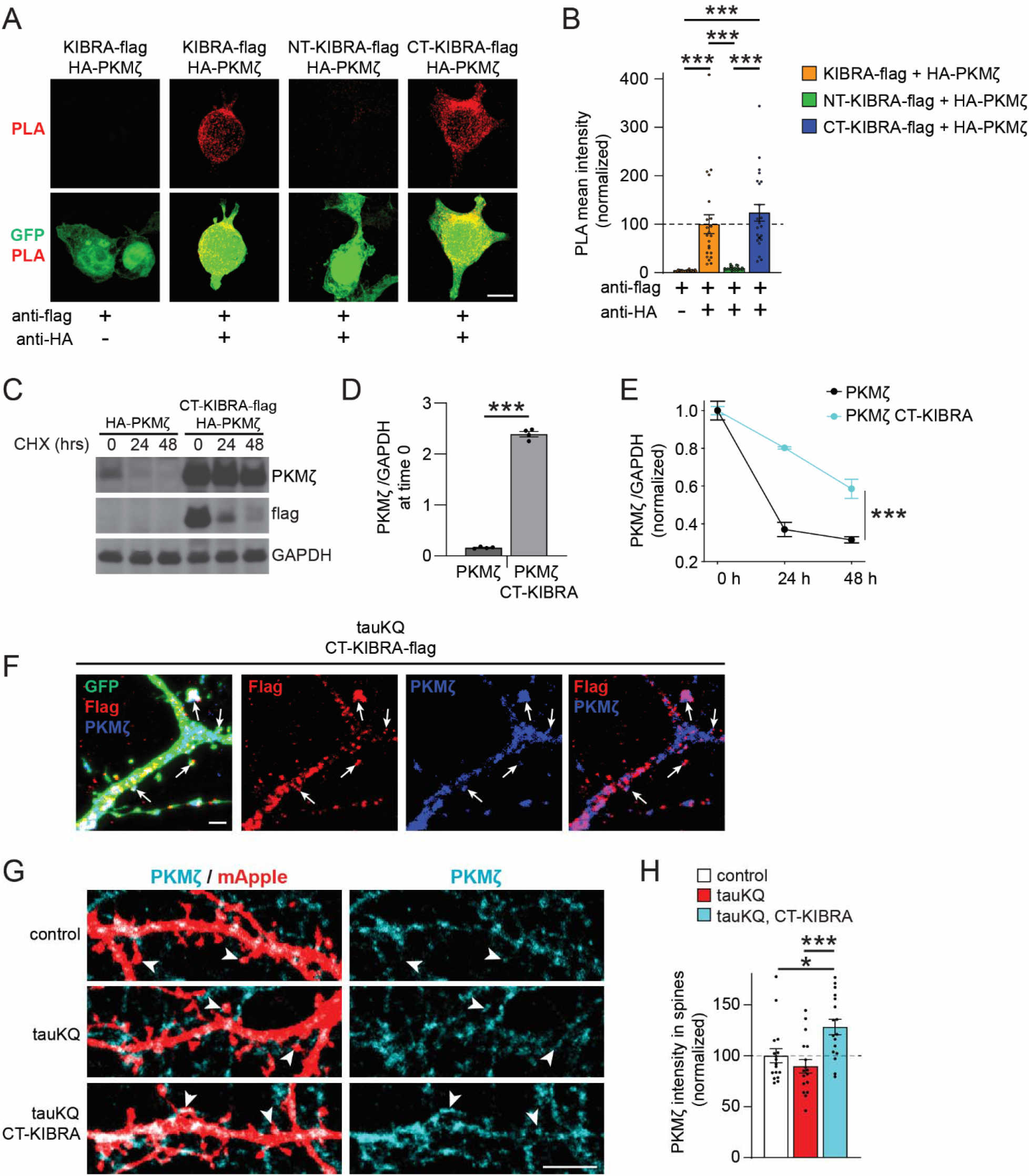
CT-KIBRA interacts with and stabilizes PKMζ at synapses in neurons with pathogenic tau. **(A)** Representative images of HEK293T cells co-expressing GFP (green) with HA-PKMζ and flag-tagged KIBRA variants. The proximity ligation assay (PLA) was applied to HEK293T cells using anti-HA and anti-flag antibodies to label only the interacting PKMζ and KIBRA variants (red) in close proximity (<40 nm). Scale bar, 10 µm. **(B)** Graph of mean PLA intensity quantification from HEK293T cells co-expressing HA-PKMζ with flag-tagged KIBRA variants normalized to PLA intensity of cells with full-length KIBRA expression (n = 15-20 cells/group; *** p < 0.001, one-way ANOVA, Bonferonni *post*-hoc analyses). See also Supplemental Figure 4. **(C-E)** HA-PKMζ and CT-KIBRA-flag expression constructs were transfected into HEK293T cells. One day later, the cells were treated with cycloheximide (CHX) to inhibit translation for 24 or 48 h. **(C)** Immunoblots of HA-PKMζ, CT-KIBRA-flag and GAPDH from HEK293T cells treated with CHX to monitor PKMζ degradation. **(D)** Quantification of PKMζ immunoblotting results normalized to GAPDH at time 0 before CHX treatment (n = 4 replicates/group; ***p < 0.001, unpaired Student’s *t*-test). **(E)** Graph showing HA-PKMζ stability in HEK293T cells with or without CT-KIBRA-flag co-expression. HA-PKMζ immunoreactivity at 0, 24h and 48h time points during the CHX treatment was normalized to the basal levels at time 0 before CHX application (n = 4 replicates/group; ***p < 0.001, two-way ANOVA, Bonferonni *post*-hoc analyses). **(F)** Co-immunolabeling of endogenous PKMζ (blue) and CT-KIBRA-flag (red) expressed in cultured hippocampal neurons with GFP (green) and tauKQ. CT-KIBRA co-localized with PKMζ in postsynaptic spines (arrows) and within dendrites. Scale bar, 2 µm. **(G)** Representative images of PKMζ immunolabeling (cyan) in dendrites from mApple expressing (red) cultured hippocampal neurons with or without tauKQ and CT-KIBRA expression. PKMζ immunoreactivity was assessed in spines (arrowheads). Scale bar, 5 µm. **(H)** Graph of mean PKMζ integrated intensity quantified in postsynaptic spines normalized to control neurons without tauKQ expression (n = 17 cells/group; * p < 0.05, *** p < 0.001, one-way ANOVA, Bonferonni *post*-hoc analyses). Values are given as means ± SEM.

We next asked whether KIBRA stabilizes PKMζ in postsynaptic spines. Flag-tagged CT-KIBRA co-localized with endogenous PKMζ within postsynaptic spines and throughout the dendrites of cultured hippocampal neurons expressing tauKQ (Figure 5F). To monitor the effect of CT-KIBRA on postsynaptic PKMζ levels, cultured hippocampal neurons were co-transfected with mApple and tauKQ with or without CT-KIBRA and immunostained for PKMζ. There was a significant increase in PKMζ in postsynaptic spines of tauKQ-expressing neurons with CT-KIBRA compared to neurons with tauKQ alone and control neurons (Figure 5, G and H). These data support that the CT-KIBRA and PKMζ interaction stabilizes and enhances PKMζ levels in postsynaptic spines of neurons with pathogenic tau.

### CT-KIBRA protects against PKM**ζ** downregulation associated with plasticity and memory impairments

A persistent increase in PKMζ levels occurs in rodent neurons after LTP induction and learning (23, 47–50). In cultured neurons expressing mApple, we found that PKMζ levels were increased within spines 24 h after chemical LTP induction compared to neurons that were not stimulated (Figure 6, A and B), in agreement with previous findings in cultured neurons (50). This enhancement of PKMζ was blocked in neurons expressing tauKQ, whereas co-expression of CT-KIBRA increased PKMζ immunostaining in spines under both unstimulated and chemical LTP conditions (Figure 6, A and B). These findings support that pathogenic acetylated tau blocks the increase in PKMζ following LTP induction, which can be rescued by CT-KIBRA mediated stabilization of PKMζ in spines.

**Figure 6.**
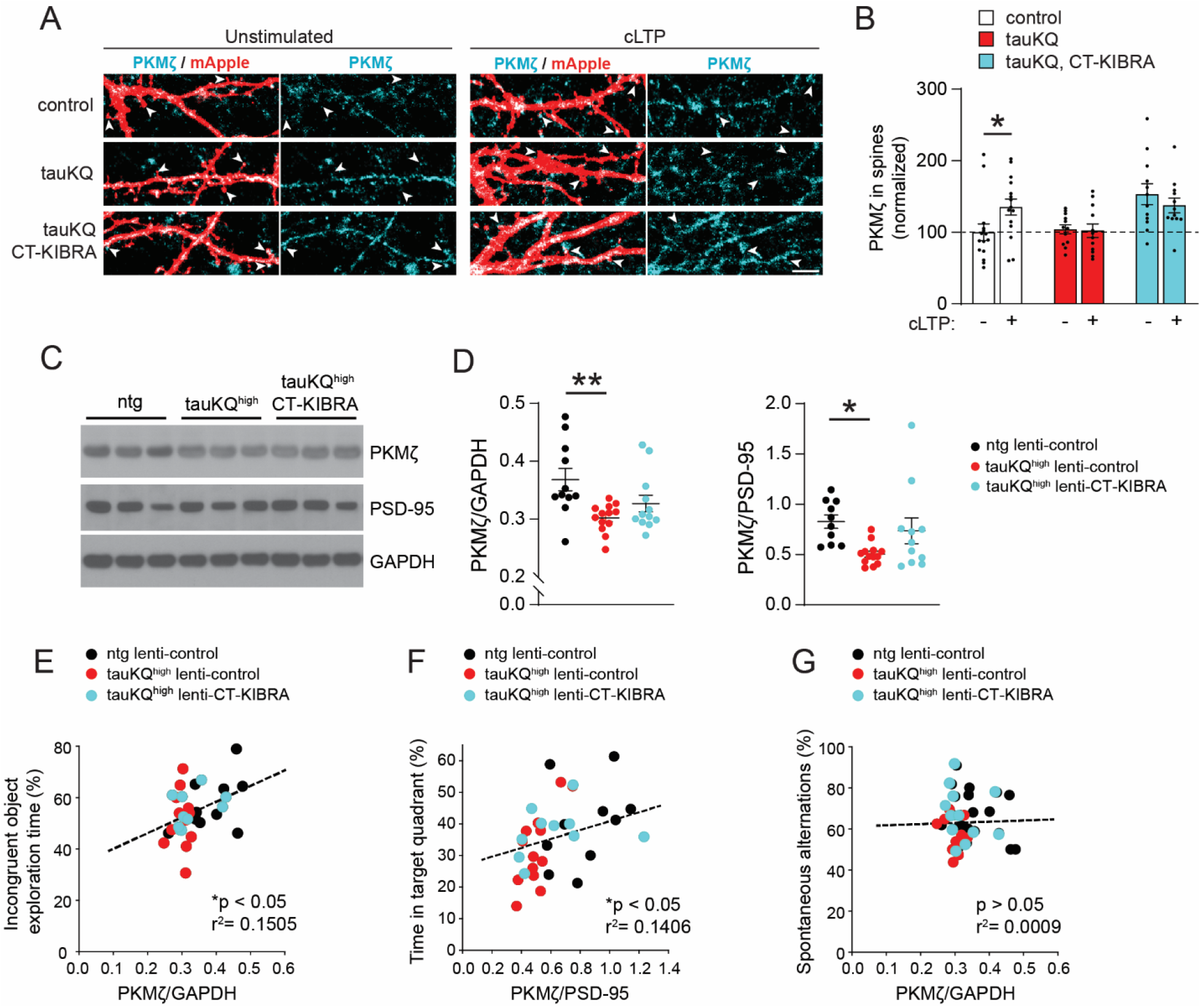
CT-KIBRA-induced resilience to tau-mediated synaptic and memory deficits is associated with higher PKMζ levels. **(A)** Representative images of dendritic spines on cultured hippocampal neurons expressing mApple (red) with or without tauKQ and CT-KIBRA. Immunolabeling of endogenous PKMζ (cyan) was monitored in spines on neurons that were unstimulated or subjected to cLTP treatment (arrowheads). Scale bar, 5 µm. **(B)** Quantification of PKMζ immunoreactivity in postsynaptic spines with or without cLTP induction normalized to postsynaptic PKMζ levels in control unstimulated neurons (n = 12-15 cells/group; * p < 0.05, unpaired Student’s *t*-test). **(C)** Representative immunoblots of PKMζ, PSD-95, and GAPDH from hippocampal homogenates of three individual ntg lenti-control, taukQ^high^ lenti-control and taukQ^high^ lenti-CT-KIBRA mice. **(D)** Quantification of total PKMζ levels in hippocampal homogenates normalized to GAPDH and to PSD-95 levels. Compared to ntg lenti-control mice, PKMζ levels were not significantly different in tauKQ^high^ lenti-CT-KIBRA mice, but they were significantly reduced in tauKQ^high^ lenti control mice (n = 10-13 mice/group; * p < 0.05, ** p < 0.01, one-way ANOVA, Bonferonni *post*- hoc analyses). Values are given as means ± SEM. See also Supplemental Figure 5. **(E)** Pearson correlation analyses between PKMζ levels relative to GAPDH analyzed from hippocampal homogenates and the percent time each mouse spent exploring the incongruent object in the object-context discrimination test of pattern separation memory (n = 10-12 mice/group). **(F)** Pearson correlation analyses of the proportion of time each mouse spent in the target quadrant of the MWM during the 7-day probe test and the corresponding PKMζ levels in hippocampus relative to PSD-95 (n = 10-13 mice/group). **(G)** Pearson correlation analyses showed no significant relationship between the percent of spontaneous alternations made in the Y-maze test and total PKMζ levels in hippocampus (n = 11-13 mice/group).

Hippocampal PKMζ levels detected by immunoblotting were significantly reduced in 12–14- month-old tauKQ^high^ mice compared to littermate controls, and PKMζ normalized to PSD-95, another postsynaptic protein, was also reduced in tauKQ^high^ mice (Supplemental Figure 5, A and B). We further assessed the impact of tauKQ on PKMζ levels in the context of learning and memory. PKMζ was quantified in hippocampal lysates from ntg lenti-control, tauKQ^high^ lenti-control, and tauKQ^high^ lenti-CT-KIBRA mice that completed hippocampus-dependent learning and memory tests (Figure 3). Total PKMζ levels in tauKQ^high^ lenti-control mice remained significantly lower than ntg mice after learning whether or not PKMζ was normalized to PSD-95, whereas tauKQ^high^ mice with CT-KIBRA showed a recovery of PKMζ levels (Figure 6, C and D). PICK1 was evaluated in the same mice, but the PICK1 levels were not significantly different in tauKQ^high^ lenti-control mice compared to ntg lenti-control mice (Supplemental Figure 5, C and D). Experiments using acute PKMζ knockdown in hippocampus suggest that PKMζ plays a role in the maintenance of hippocampal dependent long-term memory (49, 51), and the magnitude of PKMζ enhancement after learning correlates with the capacity for long-term memory in mice (49). Consistent with these studies we found a significant correlation between hippocampal PKMζ levels and long-term memory among the ntg lenti-control, tauKQ^high^ lenti-control, and tauKQ^high^ lenti-CT-KIBRA mice. Higher levels of PKMζ were associated with better memory retention in both the object-context discrimination test and the MWM (Figure 6, E and F). PKMζ levels did not correlate with the performance of the mice in the Y-maze test of short-term working memory (Figure 6G). These findings are consistent with the role of PKMζ in long-term as opposed to short-term memory. Given that we found a recovery of short-term memory in tauKQ^high^ lenti-CT-KIBRA mice that did not correlate with PKMζ levels, CT-KIBRA may also enhance synapse function and other cognitive skills through additional mechanisms not yet identified.

### The CT-KIBRA and PKM**ζ** interaction drives LTP repair in neurons with pathogenic tau

To investigate the mechanistic relationship between CT-KIBRA and PKMζ in plasticity, we examined whether elevated PKMζ levels are required for CT-KIBRA to restore AMPAR trafficking at synapses during LTP. An antisense oligodeoxynucleotide (PKMζ-antisense) was chosen to acutely block PKMζ expression in neurons to avoid compensation by other PKC isoforms (23), and because the ζ inhibitory peptide (ZIP) is not a specific inhibitor of PKMζ (52, 53). Cultured hippocampal neurons co-expressing tauKQ and CT-KIBRA were treated with PKMζ-antisense or a scrambled control 1 h before induction and throughout the expression of chemical LTP. The CT-KIBRA-mediated rescue of activity-induced postsynaptic AMPAR recruitment in tauKQ neurons was blocked by PKMζ-antisense (Figure 7, A and B). We then tested whether overexpression of PKMζ in neurons with pathogenic tau was sufficient to rescue synaptic plasticity. HA-tagged PKMζ was expressed in neurons and immunolabeling with an HA antibody revealed comparable levels of HA-PKMζ localized in spines of neurons with or without tauKQ (Supplemental Figure 6, A and B). Interestingly, the overexpression of HA-tagged PKMζ was not sufficient to restore AMPAR trafficking in tauKQ neurons in the absence of CT-KIBRA (Figure 7, C and D), supporting the critical role for KIBRA in the reversal of the tau-mediated plasticity deficit.

**Figure 7.**
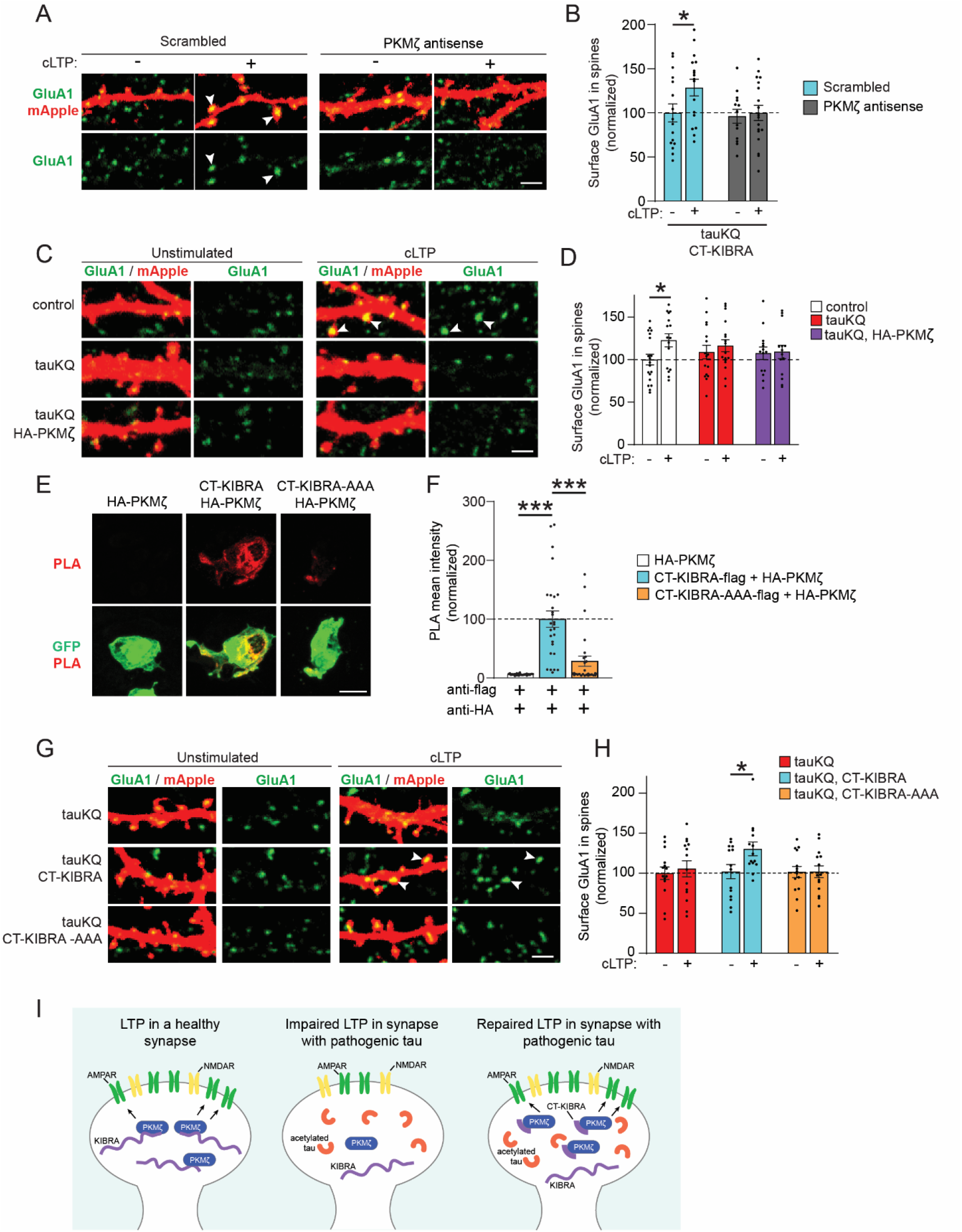
CT-KIBRA restores AMPAR trafficking during LTP by interacting with PKMζ in neurons with pathogenic tau. **(A)** Representative confocal images of surface GluA1 immunostaining (green) in spines of cultured hippocampal neurons expressing mApple (red), tauKQ, and CT-KIBRA. Unstimulated and cLTP neurons were treated with a PKMζ antisense or a scrambled control oligodeoxynucleotide. Scale bar, 2 µm. **(B)** Graph of the surface GluA1 immunofluorescence quantification in spines normalized to unstimulated neurons treated with the scrambled oligodeoxynucleotide (n = 13-18 neurons/group; * p < 0.05, unpaired Student’s *t*-test). **(C)** Representative confocal images of dendrites and spines on neurons expressing mApple (red) alone, mApple with tauKQ, or mApple with tauKQ and HA-PKMζ that were unstimulated or subjected to cLTP followed by surface GluA1 immunolabeling (green). Scale bar, 2 µm. **(D)** Graph of the mean surface GluA1 immunofluorescence quantification in spines normalized to unstimulated control neurons (n = 14-19 neurons/group; * p < 0.05, unpaired Student’s *t*-test). See also Supplemental Figure 6, A and B. **(E)** Images of HEK293T cells transfected with GFP (green) and HA-PKMζ together with CT KIBRA-flag or a CT-KIBRA-flag mutant carrying R965A, S967A and R969A mutations (CT-KIBRA-AAA-flag). Anti-flag and anti-HA antibodies were used for detection of PLA signal (red) signifying the close proximity of CT-KIBRA-flag and HA-PKMζ in the HEK293T cells. Scale bar, 10 µm. **(F)** Quantification of the mean PLA fluorescence intensity detected in HEK293T cells transfected with HA-PKMζ alone, as a control, or co-transfected with CT-KIBRA-flag constructs (n = 27-30 cells/group; *** p < 0.001, one-way ANOVA, Bonferonni *post*-hoc analyses). **(G)** Representative confocal images of dendrites and spines on neurons co-transfected with mApple (red) and tauKQ with CT-KIBRA or CT-KIBRA-AAA, which contains three alanine mutations that inhibit the interaction of CT-KIBRA with PKMζ. Scale bar, 2 µm. **(H)** Graph of the quantification of surface GluA1 immunofluorescence in spines of tauKQ expressing neurons showing that cLTP-induced postsynaptic receptor insertion is re-established by CT-KIBRA, but not CT-KIBRA-AAA. GluA1 levels were normalized to the intensity of staining in spines of unstimulated tauKQ neurons (n = 14 neurons/group; * p < 0.05, unpaired Student’s *t*-test). Values are given as means ± SEM. See also Supplemental Figure 6, C and D. **(I)** Model depicting the impact of KIBRA and PKMζ on postsynaptic AMPAR trafficking during LTP in healthy conditions (left) and in tauopathy (middle). Expression of CT-KIBRA in tauopathy neurons (right) with high pathogenic acetylated tau levels can restore postsynaptic AMPAR recruitment during plasticity, which mechanistically involves the interaction between CT-KIBRA and PKMζ.

Mutations introduced at specific residues within the C-terminal domain of KIBRA block the interaction of KIBRA with PKMζ (38). We took advantage of these mutations to test the role of the CT-KIBRA and PKMζ binding in the mechanism underlying the functional repair of synapses with pathogenic tau. We generated a CT-KIBRA-AAA mutant in which three residues, R965, S967 and R969, on the CT-KIBRA construct were changed to alanine residues (R965A, S967A and R969A), which blocks the binding of KIBRA to PKMζ (38). Using the PLA method in HEK293T cells, we confirmed that compared to wildtype CT-KIBRA, CT-KIBRA-AAA demonstrated a significantly diminished interaction with HA-PKMζ (Figure 7, E and F). In cultured neurons, the CT-KIBRA-AAA mutant was detected in dendritic spines similarly to wildtype CT-KIBRA (Supplemental Figure 6, C and D). When tested in neurons with chemical LTP, tauKQ consistently impeded postsynaptic AMPAR recruitment during plasticity, but unlike wildtype CT-KIBRA, the CT-KIBRA-AAA mutant was not sufficient to reverse the impairment (Figure 7, G and H). These findings suggest that CT-KIBRA overcomes the plasticity impairment caused by pathogenic tau by promoting synaptic resilience through its interaction with and modulation of PKMζ (Figure 7I).

## DISCUSSION

Our findings highlight the potential of KIBRA-mediated synapse repair as a novel therapeutic approach to alleviate memory loss associated with tauopathy. This targeted strategy to restore a precise functional role of KIBRA in synaptic plasticity could be leveraged to counteract the loss of KIBRA in human brain associated with cognitive impairment in tauopathy. Interestingly, CT-KIBRA improved memory performance and restored LTP in mice despite the accumulation of pathological tau in the brain, supporting that CT-KIBRA rescues mechanisms that promote synaptic resilience to tau toxicity in the brain. The restorative effect of CT-KIBRA involved its binding to and stabilization of PKMζ to promote postsynaptic AMPAR trafficking during plasticity, enabling synapses to maintain function and overcome the adverse effects of pathogenic tau.

Our study provides evidence that CT-KIBRA can improve memory when expressed in hippocampus after the onset of memory loss caused by pathogenic tau. This supports that CT-KIBRA recovers memory processes by repairing the molecular signaling at synapses required for plasticity. The enhancement of KIBRA signaling may have therapeutic potential broadly across different tauopathies because downregulated KIBRA levels were associated with elevated pathological tau in human brains across AD and Pick’s disease cases. CT-KIBRA improved synaptic plasticity and memory performance in a model of abnormal hyperacetylated tau, which is a common pathology found in most tauopathies including AD, Pick’s disease, CBD and PSP (2, 26, 54). CT-KIBRA also restored synaptic plasticity in neurons expressing human P301L tau, confirming the protective effect of CT-KIBRA against tau toxicity associated with FTD. Nonetheless, it will be important to evaluate the impact of CT-KIBRA on synapse dysfunction and cognition in other tauopathy and neurodegenerative disease models. Notably, the mechanisms by which tau drives pathophysiology in neurons in disease are complex, involving not only synapse dysfunction and synapse loss (8-10, 55, 56), but also dysregulation of axonal transport and axon initial segment function as well as axon degeneration (57–59), altered mitochondrial bioenergetics (60), nuclear transport disruption (61, 62), altered proteostasis (63), and cytoskeletal destabilization (64, 65). Given that CT-KIBRA did not affect tau levels or synapse loss in hippocampus, it likely plays a specific role in modulating synaptic plasticity.

KIBRA deficient mice have reduced PKMζ protein levels in the hippocampus, and impaired plasticity and memory loss (38, 66), which is consistent with a key role for KIBRA in PKMζ stabilization and the maintenance of synaptic plasticity. Our results show that tauKQ^high^ mice, which have reduced KIBRA at synapses (9), also had decreased PKMζ levels in hippocampus associated with impaired LTP and hippocampus-dependent memory. CT-KIBRA had a protective effect by restoring levels of PKMζ. Increased forms of pathological tau, including hyperacetylated and hyperphosphorylated tau, in human brain tissue coincided with downregulation of KIBRA, which may be driven by both tau-mediated synapse dysregulation and neuron loss. Aggregates of PKMζ have been observed in neurofibrillary tangles containing hyperphosphorylated tau in AD brain (67), but whether KIBRA is also bound to PKMζ-containing insoluble aggregates is unknown. Nevertheless, the prominent correlation between KIBRA downregulation in human brain and dementia severity suggests that KIBRA functions as a critical synaptic scaffolding protein in plasticity that can impact cognition by modulating PKMζ and other KIBRA-binding proteins at synapses.

The persistent increase in postsynaptic PKMζ levels following plasticity induction underlies the maintenance of LTP (47), and we found that this activity-dependent increase in PKMζ was blocked in neurons with pathogenic tau. Interestingly, CT-KIBRA expression enhanced the PKMζ levels in spines of tauKQ neurons with or without chemical LTP induction; yet, CT-KIBRA only increased postsynaptic AMPAR trafficking following chemical LTP without altering AMPARs in unstimulated neurons. These findings indicate that CT-KIBRA restores potentiation at synapses both by elevating PKMζ levels and by enabling an additional activity-dependent signal, which remains to be determined, that drives the AMPAR recruitment after LTP induction. Surprisingly, PKMζ overexpression alone was incapable of rescuing plasticity in tauKQ neurons, supporting that CT-KIBRA has a mechanistic effect during synaptic plasticity beyond enhancing PKMζ levels. Intriguing mechanisms that may be involved in the plasticity repair orchestrated by CT-KIBRA could involve the activation of PKMζ or other PKC isoforms, the phosphorylation of the C-terminus of KIBRA by PKMζ (40), or the phosphorylation of AMPARs. CT-KIBRA may also modify synapse strength through binding to other PKC isoforms implicated in synaptic plasticity regulation, such as PKCγ, α, β, and ι, which were reduced in postsynaptic fractions prepared from KIBRA knockout mice (66). While our study highlights the critical role for KIBRA in modulating synapse repair to recover memory, and showed that PKMζ overexpression alone is not sufficient to restore plasticity in neurons with pathogenic tau, a recent study found that the memory impairments caused by oligomeric amyloid β toxicity in rats were alleviated by PKMζ overexpression in hippocampus (68). Thus, enhanced PKMζ levels may be sufficient to promote synapse function under certain pathogenic conditions but not others. How KIBRA, PKMζ, and other PKC family members could be modulated to restore plasticity in neurodegenerative diseases deserves further investigation.

The KIBRA protein has multiple functional domains that engage in interactions with numerous postsynaptic proteins to modulate synapse function, and here we identified a specific KIBRA domain and interaction that can have a beneficial impact in the context of disease. The N-terminal WW domains of KIBRA bind to dendrin, and an inhibitory peptide that blocks this interaction reduced KIBRA levels and AMPARs at synapses on neurons, blocked LTP, and impaired memory in mice (36), indicating that the interaction between KIBRA and dendrin regulates synaptic KIBRA localization and plasticity. We expressed NT-KIBRA, comprised of only the WW domains, in neurons and detected it in dendritic spines, although to a lesser degree compared to full-length KIBRA, and it did not rescue LTP in neurons with pathogenic tau. This suggests that while the dendrin/KIBRA WW domain interaction is necessary for plasticity in healthy neurons, it is not sufficient to reverse the plasticity impairment caused by pathogenic tau. The binding of KIBRA to PICK1 can also modulate activity-dependent AMPAR trafficking in neurons (20). Our findings show a weakened interaction of CT-KIBRA with PICK1 compared to full-length KIBRA and a modest effect of pathogenic tau on PICK1 protein levels, but whether CT-KIBRA could influence PICK1-dependent AMPAR trafficking in neurons, or additional plasticity mechanisms, remains to be determined. In future studies it will be important to consider that the impact KIBRA has on synapse function and plasticity through its interactions with postsynaptic proteins may depend on normal or pathological conditions in the brain.

Our study also reveals compelling evidence that KIBRA levels in CSF are associated with tau biomarkers and cognition in AD. These results point toward KIBRA as a biomarker for synapse dysfunction in tauopathy that could be useful for diagnosis and staging disease progression. The higher KIBRA levels detected in CSF may be linked to the downregulation of KIBRA in the synaptic compartment in tauopathies. For example, KIBRA may be expelled from neurons with dysfunctional synapses and accumulate in the CSF, but the mechanisms underlying the changes in KIBRA levels at synapses and in CSF are unclear. Our findings are consistent with prior studies showing higher SNAP-25, synaptotagmin-1, GAP-43, and neurogranin levels in CSF are associated with cognitive impairment in AD (30–32), which may be linked to synapse degeneration in the brain (69). To advance clinical application and potentially identify individuals that could benefit from a KIBRA therapeutic, future studies should focus on evaluating KIBRA as a biomarker of synapse dysfunction and cognitive decline together with engineering a CT-KIBRA-based therapeutic that can be delivered across the blood-brain barrier into the brain. CT-KIBRA-mediated synapse repair could be a valuable approach in combination with pathology-modifying strategies designed to slow progression of cognitive decline by reducing tau levels or clearing toxic forms of tau from the brain which are being tested in ongoing clinical trials (70, 71). Our work supports a KIBRA-based synapse repair therapy that could promote the recovery of cognition in tauopathy with treatment by boosting the resilience of synapse function.

## ACKNOWLEDGEMENTS

We thank Dr. Lisa Ellerby, Dr. Sofiya Galkina, Jeff Simms and Dr. T. Michael Gill for support with mouse behavior tests, Stella Breslin and Harris Ingle for microscopy support, Dr. Yungui Zhou for molecular cloning support, Mary Redwine, Athena Schlereth, and Argentina Lario Lago for administrative support. This work was supported by the NIH (R03AG063248 and K01AG057862 to T.E.T, R01AG054214 to L.G.), the Buck Institute Impact Circle, the BrightFocus Foundation (A2016360F to T.E.T.), the Alzheimer’s Association (AARFD-19-616386 to G.K.), and the Larry L. Hillblom Foundation (to J.H.K). K.A.P-N was supported by an NIH T32 training grant (T32AG00026624). We acknowledge the support of instrumentation from the NCRR shared instrumentation grant 1S10 OD016281 (Buck Institute). Human tissue samples were provided by the Neurodegenerative Disease Brain Bank at the University of California, San Francisco, which receives funding support from NIH grants P01AG019724 and P50AG023501, the Consortium for Frontotemporal Dementia Research, and the Tau Consortium.

## AUTHOR CONTRIBUTIONS

T.E.T, G.K., K.A.P-N., K.B.C. and L.G. conceived the project and designed experiments. G.K. K.A.P-N., T.E.T, L.Y., J.H.C., I.W., and B.S. performed experiments. G.K., K.A.P-N., T.E.T, K.B.C., L.Y., B.S. and R.S. analyzed data. T.C.S., A.L.N, S.S., L.T.G, W.W.S., and J.H.K. provided experimental reagents. J.H.C., I.W., H.C., S.S., Y.L., and D.L. provided technical support. T.E.T, G.K. K.A.P-N., and K.B.C. wrote the manuscript.

## CONFLICT OF INTEREST STATEMENT

The authors declare no competing interests.

## SUPPLEMENTAL FIGURES AND TABLES

**Supplemental Figure 1.**
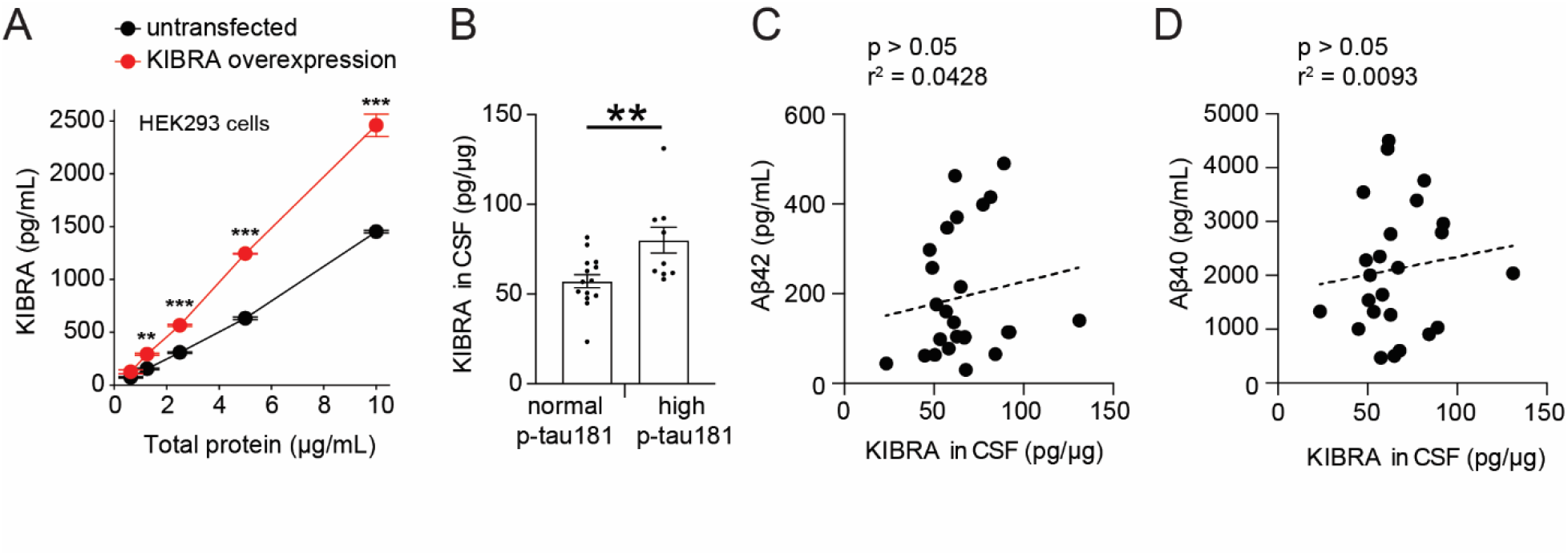
Human KIBRA detection by ELISA and CSF KIBRA relative to phosphorylated tau and Aβ related to Figure 1. **(A)** ELISA-based detection of KIBRA levels from lysates of HEK293 cells transfected with a human KIBRA construct (red) and untransfected control HEK293 cells (black). HEK293 cells with KIBRA overexpression had significantly higher KIBRA levels detected by ELISA at the different concentrations of total protein tested (n = 2 cultures/group; ** p < 0.01, *** p < 0.001, two-way ANOVA, Bonferonni posŕ-hoc analyses). Values are given as means ± SD. **(B)** Quantification of the concentration of KIBRA in total protein detected in cerebrospinal fluid (CSF) of human subjects that had either normal (< 61 pg/mL) or high (> 61 pg/mL) levels of phosphorylated tau (p-taul 81) in CSF associated with Alzheimer’s disease. Values are given as means ± SEM. **(C and D)** Spearman correlation analyses show the relationship between levels of KIBRA with concentrations of **(C)** Aβ42 and **(D)** Aβ40 in CSF (n = 24 subjects).

**Supplemental Figure 2.**
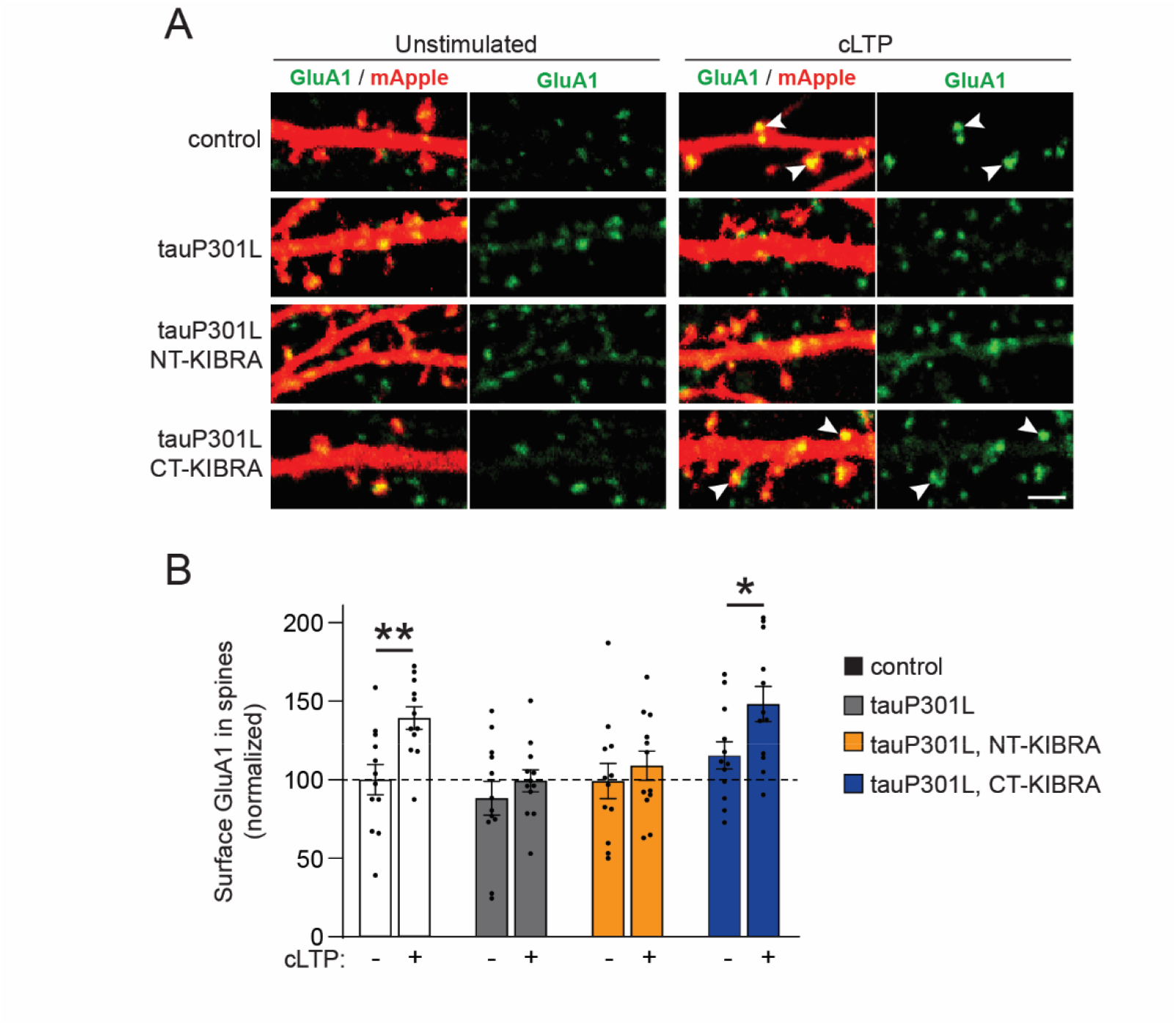
CT-KIBRA rescues AMPAR trafficking during LTP in neurons expressing human tau with the P301L mutation that causes FTD related to Figure 2. **(A)** Representative confocal images of surface GluA1 immunolabeling (green) in spines on cultured hippocampal neurons expressing mApple (red) with or without tauP301L and one of the KlBRA variants. Scale bar, 2 µm. **(B)** Quantification of surface GluA1-containing AMPARs in spines revealed that tauP301L expression in neurons inhibited chemical LTP (cLTP)­induced postsynaptic AMPAR insertion, which was restored by the co-expression of CT-KIBRA (n = 12 neurons/group; * p < 0.05, ** p < 0.01, unpaired student’s t-test). Values are given as means ± SEM.

**Supplemental Figure 3.**
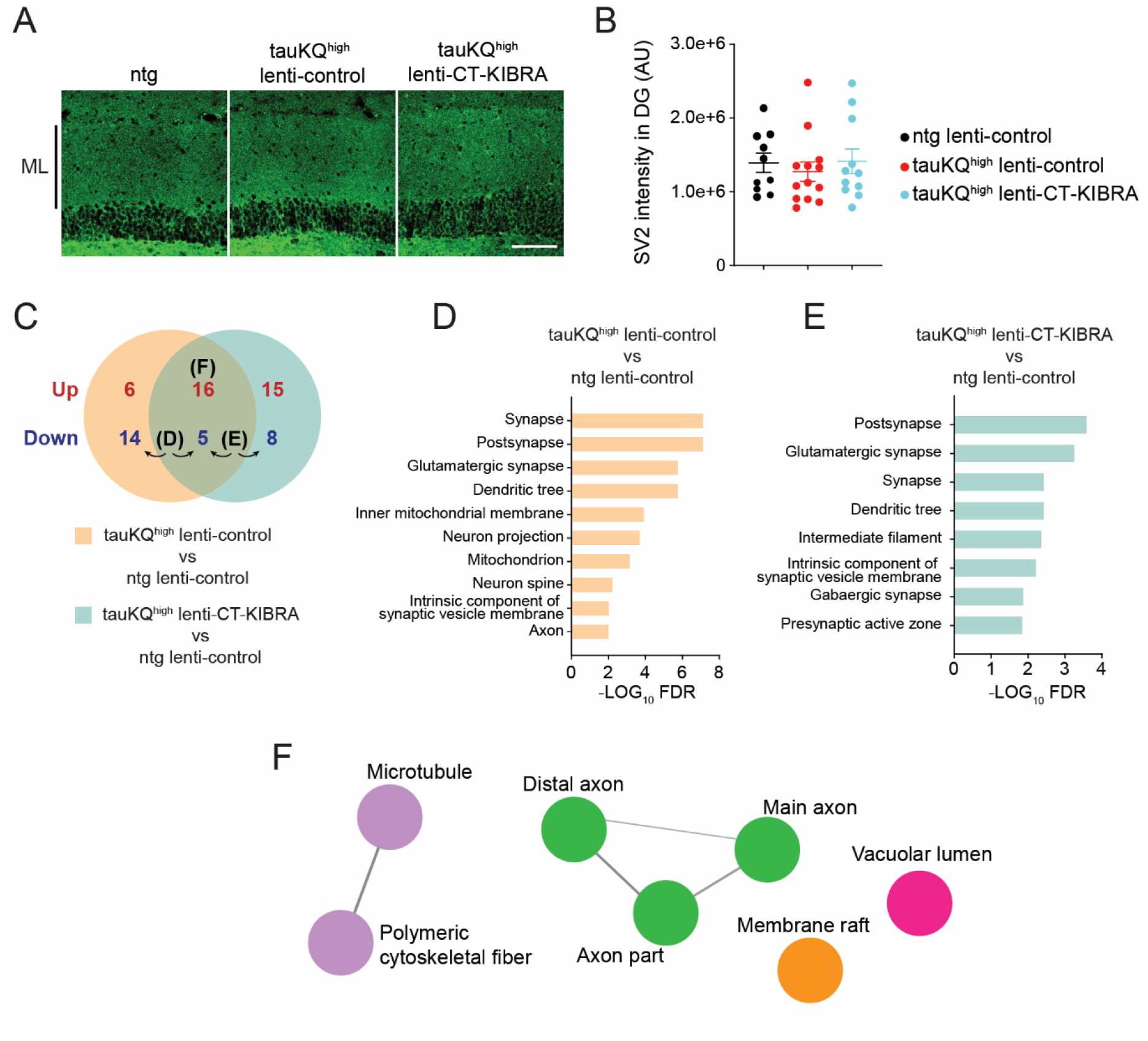
Synapse and proteomics analyses of hippocampus from tauKQ^t1iøh^ mice with or without CT-KIBRA expression related to Figure 4. **(A)** Representative confocal images of SV2 immunolabeling as a marker of synapses in the molecular layer (ML) of the dentate gyrus (DG). Scale bar, 100 µm. **(B)** Quantification of the mean integrated intensity of SV2 immunofluorescence in the ML of the DG (n = 10-13 mice/group; p > 0.05, one-way ANOVA, Bonferonni post-hoc analyses). Values are given as means ± SEM. **(C)** Venn diagram depicting the number of proteins identified that were either upregulated or downregulated in tauKQ^h’øľ^ lenti-control or tauKQ^h’øh^ lenti-CT-KIBRA compared to non-transgenic (ntg) control mice (n = 4 mice/group). See also Supplemental Table 3. **(D and E)** Gene Set Enrichment Analysis (GSEA) on the proteins identified that were downregulated in **(D)** tauKQ^Nah^ lenti-control mice and in **(E)** tauKQ^hiøh^ lenti-CT-KIBRA mice compared to controls. **(F)** ClueGO cellular component enrichment of proteins that were upregulated in hippocampus of tauKQ^h’gh^ mice with or without CT-KIBRA expression compared to ntg lenti-control mice. Node colors denote grouped networks (p < 0.0005).

**Supplemental Figure 4.**
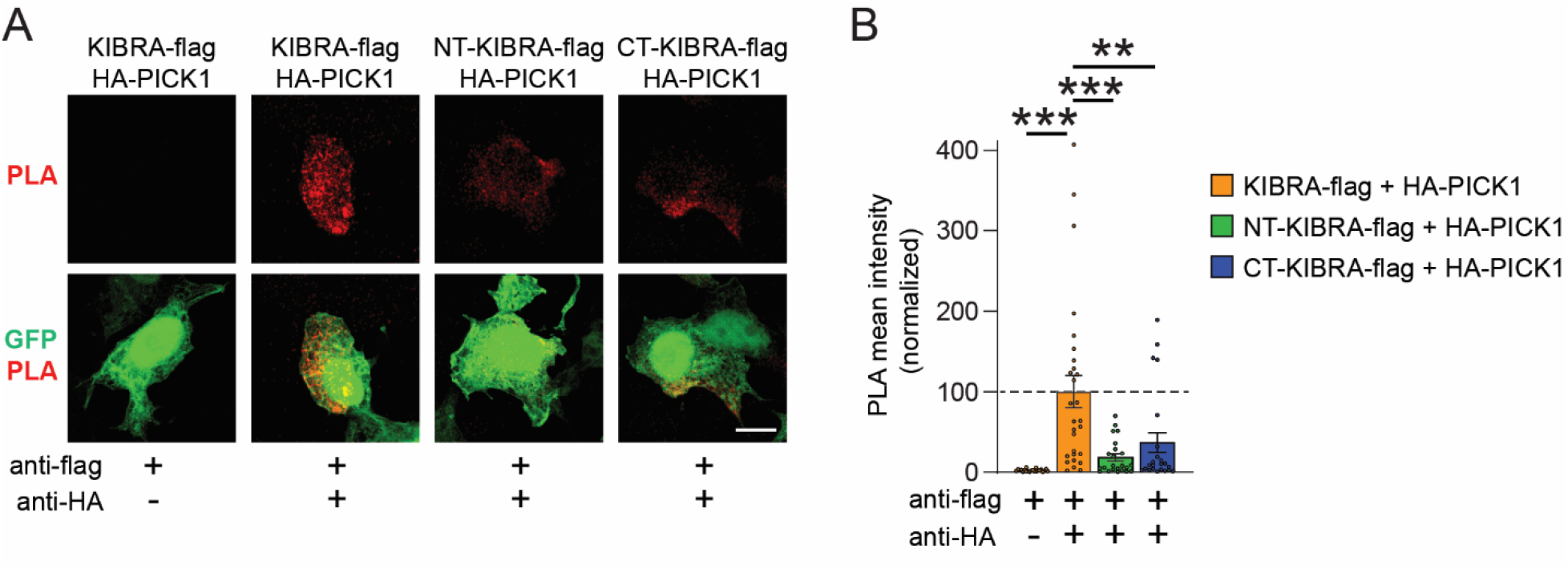
PICK1 interacts more with full-length KIBRA than the truncated KIBRA variants related to Figure 5. **(A)** HEK293 cells were transfected with GFP (green), HA-tagged PICK1 and either flag-tagged full-length KIBRA, NT-KIBRA or CT-KIBRA constructs. The proximity ligation assay (PLA) was performed with anti-flag and anti-HA antibodies to detect the PICK1 and KIBRA interacting within close proximity (red). Scale bar, 10 µm. **(B)** Graph of mean PLA intensity quantification in HEK293 cells normalized to PLA signal in cells with full-length KIBRA (n = 22-28 cells/group; *** p < 0.001, one-way ANOVA, Bonferonni post-hoc analyses). Values are given as means ± SEM.

**Supplemental Figure 5.**
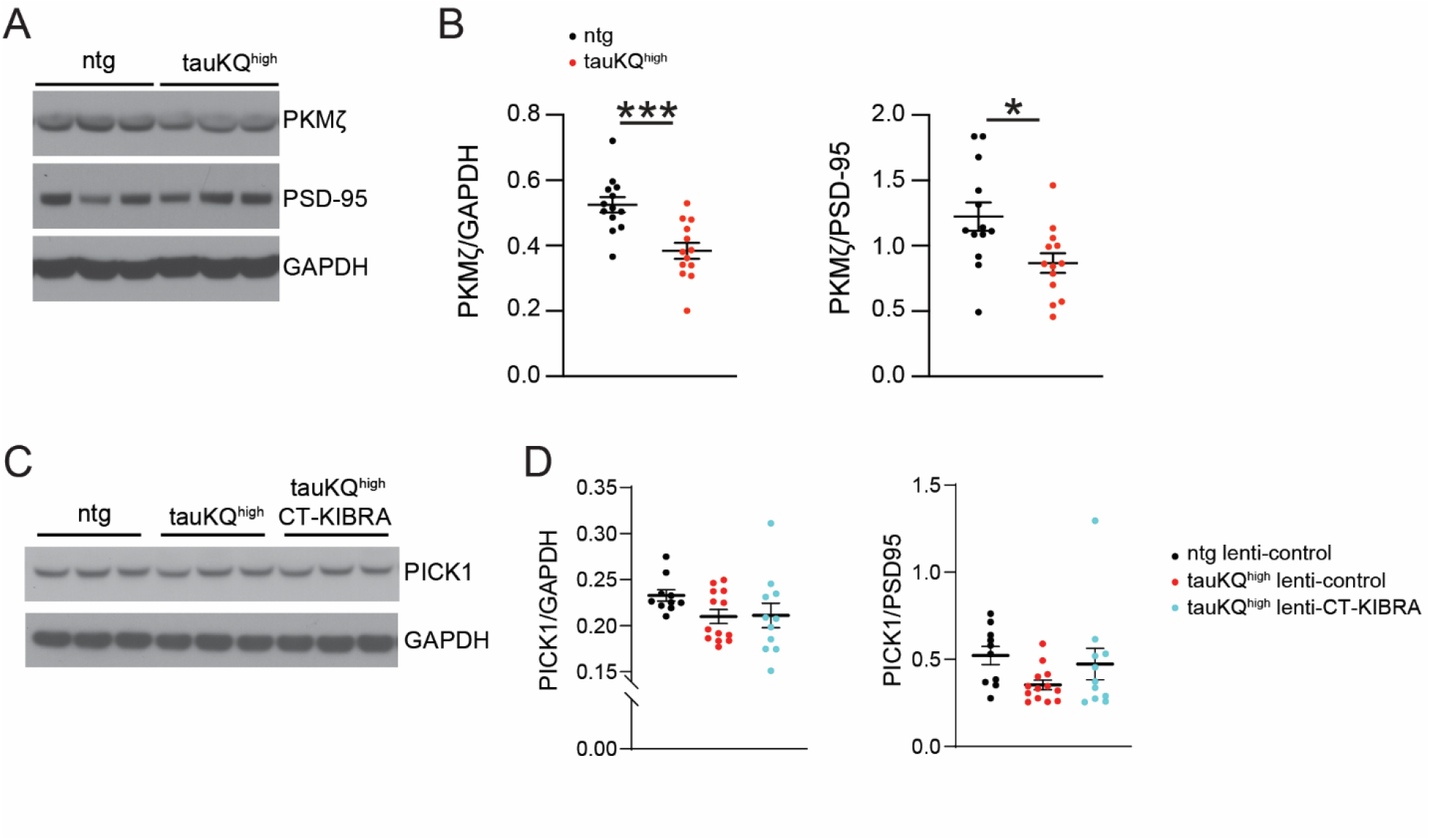
PKMζ and PICK1 levels in hippocampus of tauKQ^high^ mice related to Figure 6. **(A)** Immunoblots of PKMζ, PSD-95 and GAPDH from hippocampal lysates of three non-transgenic (ntg) and three tauKQ^h’gh^ mice that did not receive training in behavioral tests of hippocampus-dependent learning and memory. **(B)** Graphs of PKMζ levels relative to GAPDH and PSD-95 in mice that did not have training in behavioral tests (n = 13 mice/group; * p < 0.05, *** p < 0.001, unpaired student’s ŕ-test). **(C)** Immunoblots of PICK1 and GAPDH from hippocampal lysates of three mice from ntg lenti-control, tauKQ^hıgh^ lenti-control, and tauKQ^hıgh^ lenti-CT-KIBRA groups that underwent training in learning and memory tests. **(D)** Graphs of PICK1 levels in hippocampal lysates relative to GAPDH and PSD-95 (n = 10-13 mice/group; no significant differences between groups, one-way ANOVA, Bonferonni post-hoc analyses). Values are given as means ± SEM.

**Supplemental Figure 6.**
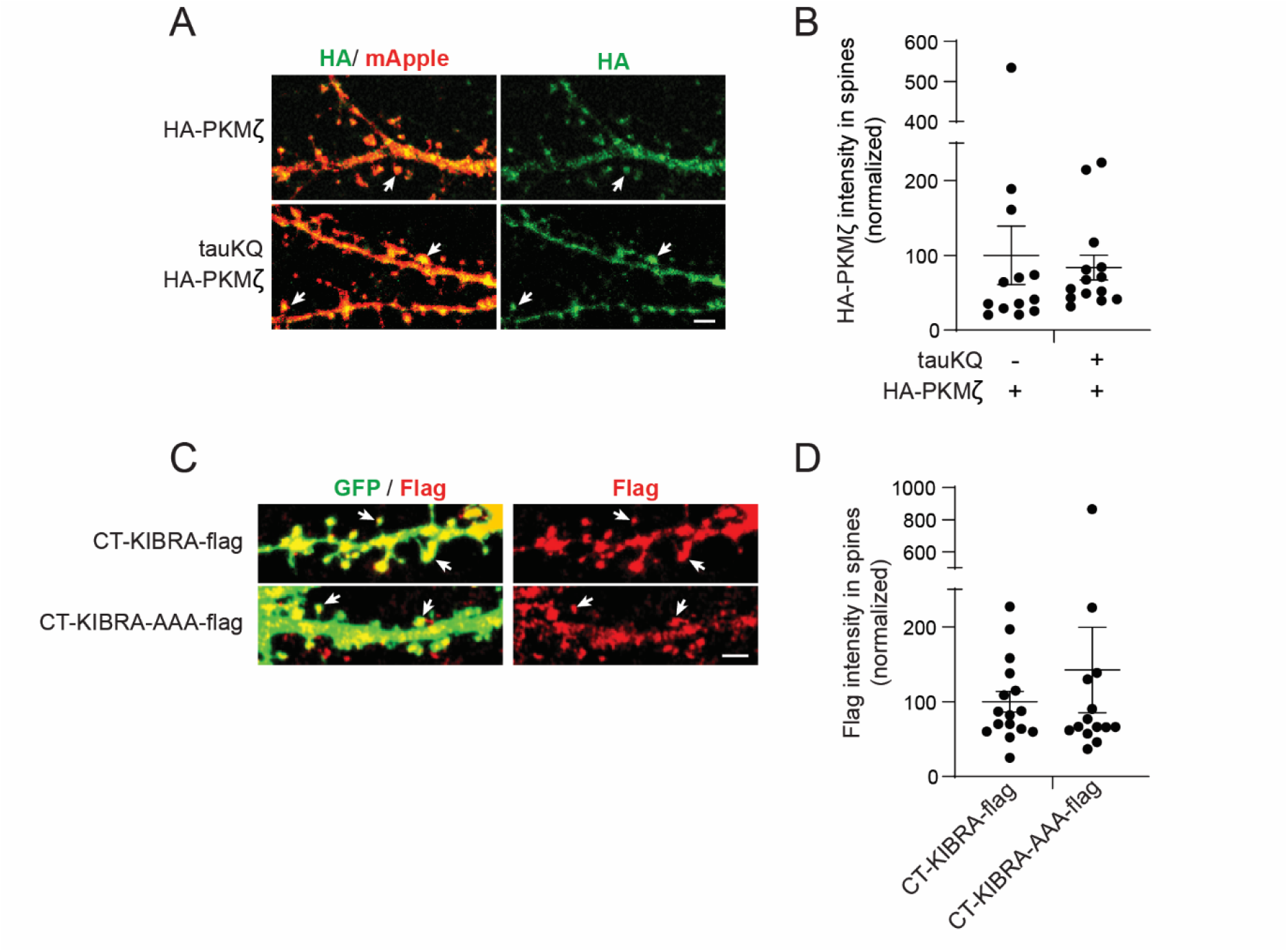
Expression of HA-PKMζ and CT-KIBRA constructs in cultured hippocampal neurons related to Figure 7. **(A)** Representative confocal images of anti-HA immunolabeling (green) of HA-tagged PKMζ overexpressed in neurons with mApple (red). HA-PKMζ was detected in spines (arrows). Scale bar, 2 µm **(B)** Quantification of the mean intensity of HA-tagged PKMζ immunolabeling in spines normalized to neurons without tauKQ co-expression (n = 13-14 neurons/group). **(C)** Representative confocal images of GFP-expressing neurons (green) with co-expression of flag-tagged CT-KIBRA or CT-KIBRA-AAA immunostained with an anti-flag antibody (red). CT-KIBRA with or without the three alanine mutations was detected in spines (arrows). Scale bar, 2 µm. **(D)** Quantification of the mean intensity of flag immunofluorescence detected in spines of neurons with CT-KIBRA-AAA compared to CT-KIBRA (n = 14-16 cells/group). Values are given as means ± SEM.

**Supplemental Table 1.** Patient information for human brain tissues used in this study.

| Case number | Sex | Age at death | Clinical | Primary Neuropath Dx | PMI (hrs)<sup>a</sup> | ADNC<sup>b</sup> | CDR | CDRsum |
| --- | --- | --- | --- | --- | --- | --- | --- | --- |
| P2323.10 | M | 70 | bvFTD | Pick's | 10.2 | - | x | x |
| P2494 | M | 72 | bvFTD | Pick's | 24.3 | Not ADNC | x | x |
| P2533 | F | 76 | bvFTD | Pick's | 25.5 | Not ADNC | 3 | 18 |
| P2682 | F | 74 | nfvPPA | Pick's | 9.8 | Not ADNC | 1 | 5.5 |
| P2839 | M | 86 | AD, Vascular | Pick's | 14.8 | Low | 3 | 18 |
| P3043 | F | 76 | nfvPPA | Pick's | 5.9 | Low | x | x |
| P3052 | M | 78 | svPPA (right) | Pick's | 7.5 | Not ADNC | 3 | 17 |
| P2345.11 | F | 86 | control | none | 7.8 | Low | x | x |
| P2577 | M | 94 | control | none | 20 | Low | x | x |
| P2594 | F | 86 | control | none | 6.4 | Not ADNC | 3 | 13 |
| P2850 | F | 92 | control | none | 8 | Not ADNC | 1 | 5.5 |
| P2921 | F | 82 | control | none | 9.6 | Low | x | x |
| P3045 | F | 84 | control | none | 7.7 | Low | 0 | 0 |
| P2412.12 | M | 88 | AD | AD | 10.8 | High | x | x |
| P2484 | M | 88 | AD | AD | 9.8 | High | x | x |
| P2508 | M | 82 | AD | AD | 6.7 | High | 3 | 16 |
| P2554 | M | 85 | AD | AD | 11.2 | High | 1 | 6 |
| P2592 | M | 86 | AD | AD | 8.6 | High | 2 | 12 |
| P2612 | M | 82 | AD | AD | 9 | High | 3 | 18 |
| P2823 | F | 89 | AD | AD | 15.8 | High | 3 | 18 |

<sup>a</sup>PMI = Post-mortem interval
<sup>b</sup>ADNC = Alzheimer's disease neuropathological change

**Supplemental Table 2.** Patient information for the cerebrospinal fluid (CSF) used in this study.

| Case number | Age | Sex | Diagnosis | MMSE<sup>a</sup> |
| --- | --- | --- | --- | --- |
| 2720 | 80 | F | normal | 28 |
| 5064 | 58 | M | normal | x |
| 7261 | 70 | M | normal | 29 |
| 7841 | 69 | F | normal | 30 |
| 8843 | 70 | M | normal | 29 |
| 12955 | 66 | M | AD | 18 |
| 13690 | 62 | M | AD | 24 |
| 14744 | 65 | M | AD | 26 |
| 15035 | 69 | F | AD | 30 |
| 15313 | 74 | F | normal | 29 |
| 15468 | 57 | M | AD | 23 |
| 15508 | 60 | F | normal | x |
| 15590 | 68 | F | AD | 21 |
| 15791 | 56 | F | AD | 25 |
| 16304 | 60 | F | AD | 8 |
| 17412 | 63 | M | AD | 12 |
| 17615 | 53 | M | normal | x |
| 17735 | 71 | F | normal | 30 |
| 18176 | 55 | M | normal | x |
| 21514 | 68 | F | normal | 30 |
| 25992 | 54 | F | normal | 29 |
| 26161 | 80 | M | AD | 25 |
| 26234 | 72 | M | normal | 30 |
| 26323 | 55 | M | normal | 30 |
| 29442 | 69 | M | normal | 29 |

<sup>a</sup>MMSE = Mini-Mental State Examination

## METHODS

### Mice

TauKQ^high^ mice were previously generated (9), and maintained in the C57BL/6 genetic background. TauKQ^high^ mice were crossed with FVB/N mice (Jackson Laboratories) to generate tauKQ^high^ mice in an FVB/N and C57BL/6 mixed background for experiments. Behavior experiments were performed in daylight hours. Mice were housed in a pathogen-free barrier facility with a 12 h light-dark cycle and provided with ad libitum access to water and food. Female and male mice were used in all experiments.

### Antibodies

Monoclonal antibodies used included: AT270 (MN1050, Thermo Fisher Scientific, RRID: AB_223651), AT180 (MN1040, Thermo Fisher Scientific, RRID: AB_223649), HT7 (MN1000, Thermo Fisher Scientific, RRID: AB_2314654), Tau5 (AHB0042, Thermo Fisher Scientific, RRID: AB_2536235), MAb359 (from Dr. Li Gan), anti-GAPDH (MAB374, Sigma-Aldrich, RRID: AB_2107445), anti-FLAG (F1804, Sigma-Aldrich, RRID: AB_262044), AT8 (MN1020, Thermo Fisher Scientific, RRID: AB223647), anti-SV2 (University of Iowa DSHB, RRID: AB_2315387), anti-PICK1 (75-040, Antibodies Inc, RRID: AB_2164544), anti-KIBRA (sc-133374, Santa Cruz Biotechnology, RRID:AB_2216359). Polyclonal antibodies used included: anti-PKMζ (from Dr. Todd C. Sacktor), anti-PSD95 (2507, Cell Signaling Technology, RRID: AB_561221), anti-HA (H6908, Sigma-Aldrich, AB_260070), GFP (A-21311, Thermo Fisher Scientific, RRID: AB_221477), anti-GluA1 (ABN241, Millipore, RRID: AB_2721164)

### Cultured Neurons

Primary hippocampal neurons were prepared from embryonic day 22 rat brains. Neurons were plated at a density of 100,000 cells / mL on coverslips coated with poly-L-lysine (Sigma) in Neurobasal medium with B27 supplement (Gibco) and Glutamax (Invitrogen). Half the media was replaced at 1 day in vitro (DIV). At 7 DIV, half the media was replaced again and 4 µM cytosine β-D-arabinofuranoside (Sigma) was added. Neurons were transfected with plasmids using Lipofectamine 2000 (Invitrogen) at 10-11 DIV.

### Chemical LTP and Immunocytochemistry

Chemical LTP experiments were performed at 14-15 DIV in extracellular solution (ECS) containing (in mM): 125 NaCl, 5 KCl, 25 HEPES, 1 NaH_2_PO_4_, 11 Glucose, 2.5 CaCl_2_, 0.0005 tetrodotoxin, 0.1 picrotoxin, and 0.001 strychnine (pH 7.4) warmed to 37°C. Neurons were briefly washed in ECS then transferred to ECS containing 300 µM Glycine for 5 min at room temperature to induce chemical LTP while unstimulated neurons were washed with ECS. After chemical LTP induction, all neurons were washed with ECS and incubated for 10 min at 37°C. Coverslips were treated with anti-GluA1 antibody (ABN241, Sigma) in ECS for 15 min at 37°C. After antibody labeling, neurons were briefly washed with ECS then fixed in 4% paraformaldehyde (PFA). Coverslips were washed three times in PBS for 10 min then incubated in blocking solution containing PBS, 2% normal goat serum, and 0.1% Triton X-100 for 1 h. An anti-rabbit conjugated Alexa 647 (A21245, Invitrogen) secondary antibody was applied for 1 h at room temperature then the coverslips were washed three times in PBS for 10 min. For experiments using GFP-TauKQ or GFP-tauP301L, neurons were labeled with anti-GFP conjugated to Alexa 488 (A21311, Invitrogen) to enhance the GFP signal. For immunolabeling of flag-tagged KIBRA, NT-KIBRA and CT-KIBRA using the anti-flag antibody, cells were fixed in 4% PFA, washed three times with PBS for 10 min then incubated in PBS with 0.1% Triton X-100 for 5 min, washed twice in PBS for 10 min, and blocked for 10 min in PBS with 1% bovine serum albumin (BSA). Primary antibodies were added in PBS with 1% BSA for 1 h, followed by three 10 min PBS washes, incubation in fluorescently conjugated secondary antibodies in PBS with 1% BSA for 1 h and three final PBS washes for 10 min. All coverslips were mounted on glass slides in Prolong Gold (Invitrogen). Images of transfected neurons were acquired on a Zeiss LSM 700 confocal microscope. Laser power and gain settings were kept constant across neurons within an experiment. Settings were established to keep the brightest pixel intensities just below saturation, except when the morphology of dendritic spines had to be clearly defined (e.g., saturated pixels of mApple fluorescence in dendrites to detect the signal in spines). For the analyses of immunolabeling within spines, the integrated intensity of GluA1, PKMζ, HA or flag immunolabeling was measured within individual dendritic spines which were outlined based on the spine morphology defined by either the mApple or GFP expressed in neurons. The intensities were analyzed from at least 50 spines per neuron using ImageJ software. Analysis was performed blind to the experimental condition.

### Stereotaxic Injection and Virus

Flag-tagged CT-KIBRA (residues 926 to 1112) was subcloned into a lentiviral FUGW2 plasmid. The FUGW2 plasmid containing CT-KIBRA was co-transfected with Δ8.9 and VSV-G packaging plasmids in HEK293T cells using calcium phosphate transfection. The lentivirus was propagated in HEK293T cells at 37°C and was collected twice over a 48-hour period. The collected media containing the lentivirus was purified using sucrose gradient ultracentrifugation. The purified lentivirus was resuspended in sterile PBS and stored at -80°C. Lentivirus titer was estimated using a p24 Rapid Titer Kit (Takara Bio USA, Inc). Mice were anesthetized by isoflurane inhalation, and a stereotax was used to position the lentivirus injection directly into the mouse hippocampus using the following coordinates from bregma: anterior-posterior: -2.0, medial lateral ±1.5, dorsal-ventral -1.8. The lentivirus was injected into the hippocampus at a rate of 0.5 uL/min. Mice received analgesic treatment with buprenorphine with one dose at the start of the surgery and two additional doses within 24 h after surgery.

### Acute Slice Preparation

Mouse brains were quickly dissected into cold sucrose dissection solution containing (in mM): 210 sucrose, 2.5 KCl, 1.25 NaH2PO4, 25 NaHCO3, 7 glucose, 2 MgSO4, and 0.5 CaCl2 (perfused with 95% O2, 5% CO2 with ph ∼7.4). Horizontal slices were cut at 400 µm thickness (VT1000S Vibratome, Leica) and incubated for 30 min in artificial ACSF at 35°C containing (in mM): 119 NaCl, 2.5 KCl, 26.2 NaHCO3, 1 NaH2PO4, 11 Glucose, 1.3 MgSO4 (gassed with 95% O2, 5% CO2 ph ∼7.4). After recovery, slices were kept at room temperature with continuous oxygen bubbling.

### Electrophysiology

Extracellular field recordings were performed from acute horizontal brain slices. Recording electrodes were placed in the molecular layer of the dentate gyrus. Slices were placed in recording chambers perfused with oxygenated ACSF solution heated to 30°C. Recording electrodes (∼3 megaohms resistance) were filled with ACSF and lowered 50 µm into the dorsal blade of the molecular layer of the dentate gyrus. A bipolar tungsten electrode (FHC) was positioned ∼150 µm away from the recording electrode to stimulate the perforant pathway inputs to the dentate gyrus. Stimulus pulses were elicited at an intensity range from 0.25 µA – 25 µA every 30 s with a 0.5 ms stimulus duration using a Model 2100 Isolated Pulse Stimulate (A-M Systems) to acquire the maximal fEPSP slope. The stimulus intensity was adjusted to 30% of the maximal fEPSP slope to record the baseline for LTP recordings. LTP recordings were performed in the presence of 100 µM picrotoxin (Sigma). Baseline fEPSPs were recorded for 20 min. Following the baseline, the stimulus intensity was increased to 60% of the maximal fEPSP slope for the theta burst stimulation (TBS) only. TBS consisted of 10 theta bursts applied every 15 s and each theta burst contained 10 bursts (4 pulses, 100 Hz) every 200 ms. After TBS the stimulus intensity was returned to the same level as during the baseline LTP recordings and fEPSPs were recorded for an additional 60 min. The fEPSP slope was normalized to baseline LTP responses. Recordings were acquired with WinLTP software (version 1.11b, University of Bristol) using a Multiclamp 700B amplifier (Molecular Devices). Recordings and analyses were performed blind to genotype.

### Behavioral Tests

For the object-context discrimination test, mice were habituated in the testing room while in their home cage for 30 min before the start of testing. During the sample phase, mice explored two similar, but distinct contexts in two different 10 min sessions that were 30 min apart. Context 1 was a white box that was cleaned with 70% ethanol. Context 2 was a white box with black and white checkered wallpaper on the 4 walls that was cleaned with 1% acetic acid. Context 1 contained two identical blue cylindrical containers (X_1_ and X_2_), and Context 2 contained two identical T25 flasks filled with yellow sand (Y_1_ and Y_2_). The mice were returned to their home cages for 4 h after the sample phase. For the test phase, one of the X objects in Context 1 was replaced with an incongruent Y object, and one of the Y objects in Context 2 was replaced with an incongruent X object. Half of the mice, balanced by genotype and sex, were tested in Context 1 and the other half were tested in Context 2. The amount of time mice spent exploring the congruent and incongruent objects during a 10-min session was calculated. The time spent exploring was established by monitoring the time the mouse’s nose was directed toward the object and within approximately 2 cm from it. Videos of the sessions were manually scored by an experimenter who was blind to mouse genotype and treatment.

Y-maze arena was constructed with plexiglass having three arms of equal length (20 cm) at equal angles. Mice were placed in the center and allowed to freely explore the arena for 5 min and recorded by video (Noldus). Each mouse was manually scored for the sequence and number of arm entries in the 5-min period and the experimenter was blind to the genotype and treatment of each mouse. An arm entry was defined as having all four paws within one arm. The percentage of spontaneous alternations for each mouse was calculated as the number of alternations divided by the total possible number of alternations in the 5-min session. An alternation is defined as the mouse having entered all three arms in succession without revisiting a previously entered arm.

The Morris water maze used a pool with a diameter of 120 cm and filled with water at a temperature 22 ± 1°C made opaque by the addition of white tempera paint. Visual cues were placed around the pool. The experimenter was blind to the genotype and treatment of each mouse. Mice were pretrained for one day with 4 trials to find a submerged hidden square platform (14 cm x 14 cm) that was 1.5 cm below the water surface located in the center of a rectangular channel. The day after pretraining, each mouse performed 4 days of hidden platform training with 2 trials per day where they had 60 s to find and sit independently on the platform for 10 s. Upon completion of hidden platform training, the platform was removed from the pool and mice were tested in a single probe trial for 60 s on subsequent days. Following probe trials, all mice were tested in cued platform training. Each mouse was recorded and tracked during hidden platform training, probe trials, and cued platform training using EthoVision XT software (Noldus).

### Western Blot

A dounce homogenizer was used to homogenize human brain tissue samples in RIPA buffer containing 50 mM Tris, pH 7.5, 150 mM NaCl, 0.5% Nonidet P-40, 1 mM EDTA, 0.5% sodium deoxycholate, 0.1% SDS, 1 mM phenylmethyl sulfonyl fluoride, protease inhibitor cocktail (Sigma), phosphatase inhibitor cocktail (Sigma), 5 mM nicotinamide (Sigma) and 1 μM trichostatin A (Sigma). Homogenized tissue was sonicated 20 times with 1 sec pulses and then centrifuged using a SW55 Ti rotor (Beckman Coulter) at 42,700 rpm for 15 min at 4°C. Mouse tissue was homogenized in RIPA buffer containing 50 mM Tris-HCl pH 7.5, 0.5% Nonidet P-40, 150 mM NaCl, 1 mM EDTA, 1 mM phenylmethyl sulfonyl fluoride, protease inhibitor cocktail (Sigma), phosphatase inhibitor cocktail 2 (Sigma) and phosphatase inhibitor cocktail 3 (Sigma). Tissue was homogenized with a hand-held homogenizer for 2 min and sonicated 10 times with 1 sec pulses. Lysates were then incubated on ice for 30 min and centrifuged at 18,000 g at 4°C for 20 min. The supernatants were collected from human or mouse brain homogenates after centrifugation and the protein concentration was measured by Bradford Assay (Bio-Rad). Equal amounts of proteins were run on a 4-12% gradient SDS-PAGE gel (Invitrogen) and the protein was transferred to a nitrocellulose membrane (GE Healthcare). The membranes were blocked with 5% nonfat dry milk in TBST at room temperature for 1 hr followed by incubation with primary antibodies in TBST with 2% nonfat dry milk overnight at 4°C. Membranes were washed with TBST and incubated with secondary antibodies in TBST with 2% nonfat dry milk at room temperature for 1 hr. After washing, chemiluminescence (Pierce) was used to detect immunolabeled proteins and ImageJ software (NIH) was used for quantification. Experiments that required quantification of immunoblots from different gels were done in parallel and 2-3 of the samples were run in duplicate on all of gels. Quantification of immunolabeling of the duplicate samples on each immunoblot was then used to normalize the immunoblot analyses across all of the immunoblots.

### Mass Spectrometry

#### Sample preparation

Mouse hippocampal tissue from 4 biological replicates for each group was homogenized in RIPA buffer (50 mM Tris-HCl pH 7.5, 0.5% Nonidet P-40, 150 mM NaCl, 1 mM EDTA, 1 mM phenylmethyl sulfonyl fluoride, protease inhibitor cocktail, and phosphatase inhibitors).

#### Proteolytic digestion

Aliquots of 150 µg protein for each tissue sample were subjected to lysis buffers containing 5% SDS and 50 mM triethylammonium bicarbonate (TEAB), at pH ∼7.55. The samples were reduced in 20 mM dithiothreitol (DTT) in 50 mM TEAB for 10 minutes at 50LJ C, subsequently cooled at room temperature for 10 minutes, and then alkylated with 40 mM iodoacetamide (IAA) in 50 mM TEAB for 30 minutes at room temperature in the dark. Samples were acidified yielding a final concentration of 1.2% phosphoric acid, resulting in a visible protein colloid. Subsequently, 90% methanol in 100 mM TEAB was added at a volume of 7 times the acidified lysate volume. Samples were vortexed until the protein colloid was thoroughly dissolved in the 90% methanol solution. The entire sample volume was spun through micro S-Trap columns (Protifi) collecting the flow-through in an Eppendorf tube (in 200 µL aliquots for 20 s at 4,000 x g), and importantly binding the samples to the S-Trap columns. Subsequently, the S-Trap columns were washed with 200 µL of 90% methanol in 100 mM TEAB (pH ∼7.1) twice for 20 s each at 4,000 x g. S-Trap columns were placed into a clean elution tube and incubated for 1 hour at 47LJ C with 125 µL of trypsin digestion buffer (in 50 mM TEAB, pH ∼8) at a 1:25 ratio (protease:protein, wt:wt). The same mixture of trypsin digestion buffer was added again for an overnight incubation at 37LJ C. Peptides were sequentially eluted from S Trap micro spin columns with 50 mM TEAB, 0.5% formic acid (FA) in water, and 50% acetonitrile (ACN) in 0.5% FA.

#### Desalting

After centrifugal evaporation, samples were resuspended in 0.2% formic acid (FA) in water and desalted with Oasis 10-mg Sorbent Cartridges (Waters, Milford, MA). Samples were then subjected to an additional centrifugal evaporation and were finally re-suspended in 0.2% FA in water with a final concentration of 1 µg/µL. One microliter of indexed Retention Time Standard (iRT, Biognosys, Schlieren, Switzerland) was added to each sample.

#### Mass spectrometry system

Briefly, samples were analyzed by reverse-phase HPLC-ESI-MS/MS using an Eksigent Ultra Plus nano-LC 2D HPLC system (Dublin, CA) with a cHiPLC system (Eksigent) which was directly connected to a quadrupole time-of-flight (QqTOF) TripleTOF 6600 mass spectrometer (SCIEX, Concord, CAN). After injection, peptide mixtures were loaded onto a C18 pre-column chip (200 µm x 0.4 mm ChromXP C18-CL chip, 3 µm, 120 Å, SCIEX) and washed at 2 µl/min for 10 min with the loading solvent (H_2_O/0.1% formic acid) for desalting. Subsequently, peptides were transferred to the 75 µm x 15 cm ChromXP C18-CL chip, 3 µm, 120 Å, (SCIEX), and eluted at a flow rate of 300 nL/min with a 3 h gradient using aqueous and acetonitrile solvent buffers.

##### Data-dependent acquisitions (for spectral library building)

For peptide and protein identifications the mass spectrometer was operated in data-dependent acquisition (DDA) mode, where the 30 most abundant precursor ions from the survey MS1 scan (250 msec) were isolated at 1 m/z resolution for collision induced dissociation tandem mass spectrometry (CID-MS/MS, 100 msec per MS/MS, ‘high sensitivity’ product ion scan mode) using the Analyst 1.7 (build 96) software with a total cycle time of 3.3 sec as previously described (72).

##### Data-independent acquisitions

For quantification, all peptide samples were analyzed by data independent acquisition (DIA, e.g., SWATH), using 64 variable-width isolation windows(73, 74). The variable window width is adjusted according to the complexity of the typical MS1 ion current observed within a certain m/z range using a DIA ‘variable window method’ algorithm (more narrow windows were chosen in ‘busy’ m/z ranges, wide windows in m/z ranges with few eluting precursor ions). DIA acquisitions produce complex MS/MS spectra, which are a composite of all the analytes within each selected Q1 m/z window. The DIA cycle time of 3.2 sec included a 250 msec precursor ion scan followed by 45 msec accumulation time for each of the 64 variable SWATH segments.

#### Mass-spectrometric data processing, quantification and bioinformatics

Mass spectrometric data-dependent acquisitions (DDA) were analyzed using the database search engine ProteinPilot (SCIEX 5.0 revision 4769) using the Paragon algorithm (5.0.0.0,4767). Using these database search engine results a MS/MS spectral library was generated in Spectronaut 14.2.200619.47784 (Biognosys). The DIA/SWATH data was processed for relative quantification comparing peptide peak areas from various different time points during the cell cycle. For the DIA/SWATH MS2 data sets quantification was based on XICs of 6-10 MS/MS fragment ions, typically y- and b-ions, matching to specific peptides present in the spectral libraries. Peptides were identified at Q< 0.01%, significantly changed proteins were accepted at a 5% FDR (q-value < 0.05). Differential expression analysis comparing ketogenic diet to wild type mouse livers was performed using a paired t-test, and p-values were corrected for multiple testing, specifically applying group wise testing corrections using the Storey method(75). Gene Ontology (GO) term enrichment analyses were performed with Gene Set Enrichment Analysis (GSEA) (76, 77) or ClueGO version 2.5.4 in Cytoscape version 3.7.1 (78, 79). Enrichment analyses were applied from GO Cellular Compartments.

### Plasmids

Flag-tagged full-length human KIBRA, NT-KIBRA and CT-KIBRA were cloned into the pCDNA 3.1 vector. The human KIBRA sequence originated from the pBabepuro-KIBRA vector (80) (Addgene #40887). For lentiviral-based expression, the flag-tagged CT-KIBRA sequence was inserted into FUGW2 (Gladstone Institutes) and packaged into virus using Δ8.9 and VSV-G plasmids. The KIBRA and CT-KIBRA sequences used were from a splice variant of KIBRA that lacks a Q residue in the KIBRA C-terminal domain (81). HA-tagged PKMζ and HA-tagged PICK1 were cloned into pcDNA 3.1 from human cDNA plasmids verified by sequencing (SinoBiological). GFP-tagged human tauKQ and tauP301L were expressed from the pEGFP-C1 plasmid (Clontech). Expression of mApple or GFP in cultured neurons was done using pGW1 mApple and pEGFP-C1 plasmids, respectively. CT-KIBRA-AAA mutant was generated by site directed mutagenesis of CT-KIBRA in pcDNA3.1 (Genscript).

### Antisense Oligonucleotide

Antisense oligonucleotides against PKMζ (C*T*C*TTGGGAAGGCAT*G*A*C) and scrambled oligonucleotides (A*A*C*AATGGGTCGTCT*C*G*G) were generated (IDT) with phosphorothioate bonds (*). For chemical LTP experiments, rat neurons were preincubated with oligonucleotides (20 µM final) for 1 h before chemical LTP, and the oligonucleotides remained in the media for the duration of the chemical LTP experiment.

### Immunohistochemistry

Mouse brain sections (30-40 mm) were used for immunohistochemistry. For AT8 immunostaining, sections were washed with PBS and heated to 95°C for 5 min in 10 mM Sodium Citrate for antigen retrieval, followed by incubation at room temperature for 20 min. Sections were washed with PBS and incubated in blocking solution (PBST with 10% normal goat serum) for 2 h at room temperature. Next, sections were incubated with primary AT8 antibody (1:1000) diluted in PBST with 3% normal goat serum overnight at 4°C. Sections were then washed with PBST, incubated with Alexa-Fluor 488 secondary antibody (1:1000) diluted in 3% normal goat serum for 2 h at room temperature and washed with PBST before mounting. Images were acquired on a digital microscope (KEYENCE) using a 20x objective lens. For SV2 immunostaining, sections were washed with PBS and non-specific binding was blocked by 10% normal goat serum in PBST for 1 h at room temperature. Next, sections were incubated with SV2 antibody (1:500) diluted in PBST with 3% normal goat serum overnight at 4°C. Sections were then washed with PBST, incubated with Alexa-Fluor 488 secondary antibody (1:1000) diluted in 3% normal goat serum for 2 h at room temperature and washed with PBST before mounting. Images were acquired on LSM 700 laser scanning confocal microscope (Zeiss) using a 20x objective lens. ImageJ software (NIH) was used to quantify immunofluorescence from both AT8 and SV2 immunostaining.

### Proximity Ligation Assay (PLA)

HEK293T cells were plated on glass coverslips coated with poly-L-lysine at a density of 80,000 cells/mL. Cells were transfected 24 h later with plasmid DNA with Lipofectamine 2000 (Invitrogen). HEK cells were fixed in 4% PFA 24 h after transfection and washed three times with PBS. Following the manufacturers protocol, coverslips were blocked in Duolink blocking solution (DUO82007, Sigma) then labeled with primary antibodies and the secondary antibodies, anti-Rabbit PLUS (DUO92002, Sigma) and anti-Mouse MINUS (DUO92004, Sigma). PLA signals were detected using the red Duolink in situ detection reagent kit (DUO92008, Sigma) and imaged on the Zeiss LSM700. Images were analyzed using ImageJ software.

### Protein Stability Assay

Plated 0.5 x 10^6^ HEK293T cells on each well of a 12-well culture plate. The cells were then transfected with HA-PKMζ or HA-PKMζ and CT-KIBRA-flag using lipofectamine 2000 (Life Technologies). A day after transfection, untreated cells (0 hr timepoint) were harvested, while the rest of the cells were treated with 5 mg/ml of cycloheximide. Treated cells were then harvested after 24 and 48 h of incubation with cycloheximide. Harvested cells were resuspended in RIPA buffer containing 50 mM Tris-HCl pH 7.5, 0.5% Nonidet P-40, 150 mM NaCl, 1 mM EDTA, 1 mM phenylmethyl sulfonyl fluoride, protease inhibitor cocktail (Sigma), phosphatase inhibitor cocktail 2 (Sigma) and phosphatase inhibitor cocktail 3 (Sigma). Resuspended cells were sonicated 10 times with 1 sec pulses. Lysates were then incubated on ice for 30 min and centrifuged at 18,000 g at 4°C for 20 min. The supernatant was collected and western blot was performed to detect PKMζ and KIBRA levels.

### Human Subjects

#### Brain tissue samples

Tissue samples were acquired from the Neurodegenerative Disease Brain Bank at the University of California San Francisco. Neuropathological diagnoses were made following consensus diagnostic criteria (82, 83), and previously described histological and immunohistochemical methods were used (84, 85). Cases were selected based on clinical and neuropathological diagnoses. Frozen human brain tissue samples were dissected from the middle temporal gyrus of control (n=6), AD (n=7) and Pick’s disease (n=7) cases.

#### CSF Samples

For in-vivo CSF KIBRA quantification, 25 older adults who were either clinically normal (n= 15) or diagnosed with Alzheimer’s dementia (n=10) were included. All participants were community-dwelling older adults who completed comprehensive neurologic and neuropsychological evaluations, as well as study partner interview at the University of California San Francisco Memory and Aging Center. Participants were reviewed and determined to be either within normative standards or meet research consensus criteria for Alzheimer’s dementia(86) at a multidisciplinary consensus conference by board-certified neurologists and neuropsychologists. All clinically normal adults evidenced no functional decline (Clinical Dementia Rating (CDR)= 0), while AD participants ranged from mild to moderate stages of functional impairment (CDR range 0.5 to 2).

CSF was collected via lumbar puncture in the morning following a 12-hour fast in sterile polypropylene tubes. Within 30 min of collection, CSF samples were centrifuged at 2000g at room temperature (20-25°C) for 5 min before being aliquoted into 500μL cryovials and stored at -80°C until analysis, following standard procedure (87).

### ELISAs

Commercially available enzyme-linked immunosorbent assay (ELISA) kits were used to measure levels of KIBRA (Wuhan Fine Biotech Co., Ltd, China), p-tau181, t-tau (INNOTEST, USA), Aβ40 and Aβ42 (ThermoFisher, USA) in CSF samples. ELISAs on CSF were performed in duplicates and averaged. Samples with % CV > 20% were excluded from analyses.

### Statistical Analyses

Differences between two means were analyzed by Student’s *t*-test, and differences between multiple means were analyzed by one-way or two-way ANOVA with Bonferroni multiple comparison *post*-hoc analyses. Repeated measures two-way ANOVA with Bonferroni multiple comparison was used to analyze input output results from field recordings of dentate gyrus and to analyze spatial learning in the MWM test. ClueGO analyses of mass spectrometry results showed networks with p-values < 0.05 with right-sided hypergeometric testing and Bonferroni adjustment. Either Spearman’s or Pearson’s correlational analyses were used to evaluate the relationship between two variables.

### Study approval

All animal procedures were completed under the supervision of the Buck Institute Research on Aging Institutional Animal Care and Use Committee. For studies on human brain tissue, prior to autopsy, patients or their surrogates provided informed consent for brain donation, in keeping with the guidelines put forth in the Declaration of Helsinki. All study procedures for human CSF samples were also conducted in accordance with the latest Declaration of Helsinki and approved by the local UCSF Institutional Review Board. Participants provided written informed consent to participate in study procedures.

## Notes

### Competing Interest Statement

The authors have declared no competing interest.

## REFERENCES

1. Alquezar C, Arya S, and Kao AW. Tau Post-translational Modifications: Dynamic Transformers of Tau Function, Degradation, and Aggregation. Frontiers in neurology. 2020;11:595532.

2. Arakhamia T, Lee CE, Carlomagno Y, Duong DM, Kundinger SR, Wang K, et al. Posttranslational Modifications Mediate the Structural Diversity of Tauopathy Strains. Cell. 2020;180(4):633–44.e12.

3. Nelson PT, Alafuzoff I, Bigio EH, Bouras C, Braak H, Cairns NJ, et al. Correlation of Alzheimer disease neuropathologic changes with cognitive status: a review of the literature. Journal of neuropathology and experimental neurology. 2012;71(5):362–81.

4. Tanner JA, and Rabinovici GD. Relationship Between Tau and Cognition in the Evolution of Alzheimer’s Disease: New Insights from Tau PET. Journal of nuclear medicine : official publication, Society of Nuclear Medicine. 2021;62(5):612–3.

5. Tai HC, Wang BY, Serrano-Pozo A, Frosch MP, Spires-Jones TL, and Hyman BT. Frequent and symmetric deposition of misfolded tau oligomers within presynaptic and postsynaptic terminals in Alzheimer’s disease. Acta neuropathologica communications. 2014;2:146.

6. Tai HC, Serrano-Pozo A, Hashimoto T, Frosch MP, Spires-Jones TL, and Hyman BT. The synaptic accumulation of hyperphosphorylated tau oligomers in Alzheimer disease is associated with dysfunction of the ubiquitin-proteasome system. The American journal of pathology. 2012;181(4):1426–35.

7. Henkins KM, Sokolow S, Miller CA, Vinters HV, Poon WW, Cornwell LB, et al. Extensive p-tau pathology and SDS-stable p-tau oligomers in Alzheimer’s cortical synapses. Brain pathology (Zurich, Switzerland). 2012;22(6):826–33.

8. Yoshiyama Y, Higuchi M, Zhang B, Huang SM, Iwata N, Saido TC, et al. Synapse loss and microglial activation precede tangles in a P301S tauopathy mouse model. Neuron. 2007;53(3):337–51.

9. Tracy TE, Sohn PD, Minami SS, Wang C, Min SW, Li Y, et al. Acetylated Tau Obstructs KIBRA-Mediated Signaling in Synaptic Plasticity and Promotes Tauopathy-Related Memory Loss. Neuron. 2016;90(2):245–60.

10. Hoover BR, Reed MN, Su J, Penrod RD, Kotilinek LA, Grant MK, et al. Tau mislocalization to dendritic spines mediates synaptic dysfunction independently of neurodegeneration. Neuron. 2010;68(6):1067–81.

11. Sydow A, Van der Jeugd A, Zheng F, Ahmed T, Balschun D, Petrova O, et al. Tau induced defects in synaptic plasticity, learning, and memory are reversible in transgenic mice after switching off the toxic Tau mutant. The Journal of neuroscience : the official journal of the Society for Neuroscience. 2011;31(7):2511–25.

12. Shipton OA, Leitz JR, Dworzak J, Acton CE, Tunbridge EM, Denk F, et al. Tau protein is required for amyloid {beta}-induced impairment of hippocampal long-term potentiation. The Journal of neuroscience : the official journal of the Society for Neuroscience. 2011;31(5):1688–92.

13. Shentu YP, Huo Y, Feng XL, Gilbert J, Zhang Q, Liuyang ZY, et al. CIP2A Causes Tau/APP Phosphorylation, Synaptopathy, and Memory Deficits in Alzheimer’s Disease. Cell reports. 2018;24(3):713–23.

14. Lasagna-Reeves CA, Castillo-Carranza DL, Sengupta U, Guerrero-Munoz MJ, Kiritoshi T, Neugebauer V, et al. Alzheimer brain-derived tau oligomers propagate pathology from endogenous tau. Scientific reports. 2012;2:700.

15. Fá M, Puzzo D, Piacentini R, Staniszewski A, Zhang H, Baltrons MA, et al. Extracellular Tau Oligomers Produce An Immediate Impairment of LTP and Memory. Scientific reports. 2016;6:19393.

16. Papassotiropoulos A, Stephan DA, Huentelman MJ, Hoerndli FJ, Craig DW, Pearson JV, et al. Common Kibra alleles are associated with human memory performance. Science (New York, NY). 2006;314(5798):475–8.

17. Corneveaux JJ, Liang WS, Reiman EM, Webster JA, Myers AJ, Zismann VL, et al. Evidence for an association between KIBRA and late-onset Alzheimer’s disease. Neurobiology of aging. 2010;31(6):901–9.

18. Burgess JD, Pedraza O, Graff-Radford NR, Hirpa M, Zou F, Miles R, et al. Association of common KIBRA variants with episodic memory and AD risk. Neurobiology of aging. 2011;32(3):557.e1-9.

19. Rodriguez-Rodriguez E, Infante J, Llorca J, Mateo I, Sanchez-Quintana C, Garcia-Gorostiaga I, et al. Age-dependent association of KIBRA genetic variation and Alzheimer’s disease risk. Neurobiology of aging. 2009;30(2):322–4.

20. Makuch L, Volk L, Anggono V, Johnson RC, Yu Y, Duning K, et al. Regulation of AMPA receptor function by the human memory-associated gene KIBRA. Neuron. 2011;71(6):1022–9.

21. Schneider A, Huentelman MJ, Kremerskothen J, Duning K, Spoelgen R, and Nikolich K. KIBRA: A New Gateway to Learning and Memory? Frontiers in aging neuroscience. 2010;2:4.

22. Zhang L, Yang S, Wennmann DO, Chen Y, Kremerskothen J, and Dong J. KIBRA: In the brain and beyond. Cellular signalling. 2014;26(7):1392–9.

23. Tsokas P, Hsieh C, Yao Y, Lesburguères E, Wallace EJC, Tcherepanov A, et al. Compensation for PKMζ in long-term potentiation and spatial long-term memory in mutant mice. eLife. 2016;5.

24. Sacktor TC, and Hell JW. The genetics of PKMζ and memory maintenance. Science signaling. 2017;10(505).

25. Ling DS, Benardo LS, and Sacktor TC. Protein kinase Mzeta enhances excitatory synaptic transmission by increasing the number of active postsynaptic AMPA receptors. Hippocampus. 2006;16(5):443–52.

26. Grinberg LT, Wang X, Wang C, Sohn PD, Theofilas P, Sidhu M, et al. Argyrophilic grain disease differs from other tauopathies by lacking tau acetylation. Acta neuropathologica. 2013;125(4):581–93.

27. Lynch CA, Walsh C, Blanco A, Moran M, Coen RF, Walsh JB, et al. The clinical dementia rating sum of box score in mild dementia. Dementia and geriatric cognitive disorders. 2006;21(1):40–3.

28. O’Bryant SE, Waring SC, Cullum CM, Hall J, Lacritz L, Massman PJ, et al. Staging dementia using Clinical Dementia Rating Scale Sum of Boxes scores: a Texas Alzheimer’s research consortium study. Archives of neurology. 2008;65(8):1091–5.

29. Lleó A, Núñez-Llaves R, Alcolea D, Chiva C, Balateu-Paños D, Colom-Cadena M, et al. Changes in Synaptic Proteins Precede Neurodegeneration Markers in Preclinical Alzheimer’s Disease Cerebrospinal Fluid. Molecular & cellular proteomics : MCP. 2019;18(3):546–60.

30. Brinkmalm A, Brinkmalm G, Honer WG, Frölich L, Hausner L, Minthon L, et al. SNAP-25 is a promising novel cerebrospinal fluid biomarker for synapse degeneration in Alzheimer’s disease. Molecular neurodegeneration. 2014;9:53.

31. Casaletto KB, Elahi FM, Bettcher BM, Neuhaus J, Bendlin BB, Asthana S, et al. Neurogranin, a synaptic protein, is associated with memory independent of Alzheimer biomarkers. Neurology. 2017;89(17):1782–8.

32. Thorsell A, Bjerke M, Gobom J, Brunhage E, Vanmechelen E, Andreasen N, et al. Neurogranin in cerebrospinal fluid as a marker of synaptic degeneration in Alzheimer’s disease. Brain research. 2010;1362:13–22.

33. Colom-Cadena M, Spires-Jones T, Zetterberg H, Blennow K, Caggiano A, DeKosky ST, et al. The clinical promise of biomarkers of synapse damage or loss in Alzheimer’s disease. Alzheimer’s research & therapy. 2020;12(1):21.

34. Rosén C, Hansson O, Blennow K, and Zetterberg H. Fluid biomarkers in Alzheimer’s disease - current concepts. Molecular neurodegeneration. 2013;8:20.

35. Vanderstichele H, De Vreese K, Blennow K, Andreasen N, Sindic C, Ivanoiu A, et al. Analytical performance and clinical utility of the INNOTEST PHOSPHO-TAU181P assay for discrimination between Alzheimer’s disease and dementia with Lewy bodies. Clinical chemistry and laboratory medicine. 2006;44(12):1472–80.

36. Ji Z, Li H, Yang Z, Huang X, Ke X, Ma S, et al. Kibra Modulates Learning and Memory via Binding to Dendrin. Cell reports. 2019;26(8):2064–77.e7.

37. Duning K, Schurek EM, Schlüter M, Bayer M, Reinhardt HC, Schwab A, et al. KIBRA modulates directional migration of podocytes. Journal of the American Society of Nephrology : JASN. 2008;19(10):1891–903.

38. Vogt-Eisele A, Krüger C, Duning K, Weber D, Spoelgen R, Pitzer C, et al. KIBRA (KIdney/BRAin protein) regulates learning and memory and stabilizes Protein kinase Mζ. Journal of neurochemistry. 2014;128(5):686–700.

39. Yoshihama Y, Sasaki K, Horikoshi Y, Suzuki A, Ohtsuka T, Hakuno F, et al. KIBRA suppresses apical exocytosis through inhibition of aPKC kinase activity in epithelial cells. Current biology : CB. 2011;21(8):705–11.

40. Büther K, Plaas C, Barnekow A, and Kremerskothen J. KIBRA is a novel substrate for protein kinase Czeta. Biochemical and biophysical research communications. 2004;317(3):703–7.

41. Lu W, Man H, Ju W, Trimble WS, MacDonald JF, and Wang YT. Activation of synaptic NMDA receptors induces membrane insertion of new AMPA receptors and LTP in cultured hippocampal neurons. Neuron. 2001;29(1):243–54.

42. Oku Y, and Huganir RL. AGAP3 and Arf6 regulate trafficking of AMPA receptors and synaptic plasticity. The Journal of neuroscience : the official journal of the Society for Neuroscience. 2013;33(31):12586–98.

43. Chen MK, Mecca AP, Naganawa M, Finnema SJ, Toyonaga T, Lin SF, et al. Assessing Synaptic Density in Alzheimer Disease With Synaptic Vesicle Glycoprotein 2A Positron Emission Tomographic Imaging. JAMA neurology. 2018;75(10):1215–24.

44. Mecca AP, Chen MK, O’Dell RS, Naganawa M, Toyonaga T, Godek TA, et al. In vivo measurement of widespread synaptic loss in Alzheimer’s disease with SV2A PET. Alzheimer’s & dementia : the journal of the Alzheimer’s Association. 2020;16(7):974–82.

45. Yoshihama Y, Hirai T, Ohtsuka T, and Chida K. KIBRA Co-localizes with protein kinase Mzeta (PKMzeta) in the mouse hippocampus. Bioscience, biotechnology, and biochemistry. 2009;73(1):147–51.

46. Söderberg O, Gullberg M, Jarvius M, Ridderstråle K, Leuchowius KJ, Jarvius J, et al. Direct observation of individual endogenous protein complexes in situ by proximity ligation. Nature methods. 2006;3(12):995–1000.

47. Osten P, Valsamis L, Harris A, and Sacktor TC. Protein synthesis-dependent formation of protein kinase Mzeta in long-term potentiation. The Journal of neuroscience : the official journal of the Society for Neuroscience. 1996;16(8):2444–51.

48. Hsieh C, Tsokas P, Grau-Perales A, Lesburguères E, Bukai J, Khanna K, et al. Persistent increases of PKMζ in memory-activated neurons trace LTP maintenance during spatial long-term memory storage. The European journal of neuroscience. 2021;54(8):6795–814.

49. Hsieh C, Tsokas P, Serrano P, Hernández AI, Tian D, Cottrell JE, et al. Persistent increased PKMζ in long-term and remote spatial memory. Neurobiology of learning and memory. 2017;138:135–44.

50. Palida SF, Butko MT, Ngo JT, Mackey MR, Gross LA, Ellisman MH, et al. PKMζ, but not PKCλ, is rapidly synthesized and degraded at the neuronal synapse. The Journal of neuroscience : the official journal of the Society for Neuroscience. 2015;35(20):7736–49.

51. Wang S, Sheng T, Ren S, Tian T, and Lu W. Distinct Roles of PKCι/λ and PKMζ in the Initiation and Maintenance of Hippocampal Long-Term Potentiation and Memory. Cell reports. 2016;16(7):1954–61.

52. Lee AM, Kanter BR, Wang D, Lim JP, Zou ME, Qiu C, et al. Prkcz null mice show normal learning and memory. Nature. 2013;493(7432):416–9.

53. Volk LJ, Bachman JL, Johnson R, Yu Y, and Huganir RL. PKM-ζ is not required for hippocampal synaptic plasticity, learning and memory. Nature. 2013;493(7432):420–3.

54. Irwin DJ, Cohen TJ, Grossman M, Arnold SE, Xie SX, Lee VM, et al. Acetylated tau, a novel pathological signature in Alzheimer’s disease and other tauopathies. Brain : a journal of neurology. 2012;135(Pt 3):807–18.

55. Lasagna-Reeves CA, Castillo-Carranza DL, Sengupta U, Clos AL, Jackson GR, and Kayed R. Tau oligomers impair memory and induce synaptic and mitochondrial dysfunction in wild-type mice. Molecular neurodegeneration. 2011;6:39.

56. McInnes J, Wierda K, Snellinx A, Bounti L, Wang YC, Stancu IC, et al. Synaptogyrin-3 Mediates Presynaptic Dysfunction Induced by Tau. Neuron. 2018.

57. Zhang B, Higuchi M, Yoshiyama Y, Ishihara T, Forman MS, Martinez D, et al. Retarded axonal transport of R406W mutant tau in transgenic mice with a neurodegenerative tauopathy. The Journal of neuroscience : the official journal of the Society for Neuroscience. 2004;24(19):4657–67.

58. Ishihara T, Hong M, Zhang B, Nakagawa Y, Lee MK, Trojanowski JQ, et al. Age dependent emergence and progression of a tauopathy in transgenic mice overexpressing the shortest human tau isoform. Neuron. 1999;24(3):751–62.

59. Sohn PD, Huang CT, Yan R, Fan L, Tracy TE, Camargo CM, et al. Pathogenic Tau Impairs Axon Initial Segment Plasticity and Excitability Homeostasis. Neuron. 2019.

60. Tracy TE, Madero-Pérez J, Swaney DL, Chang TS, Moritz M, Konrad C, et al. Tau interactome maps synaptic and mitochondrial processes associated with neurodegeneration. Cell. 2022;185(4):712–28.e14.

61. Eftekharzadeh B, Daigle JG, Kapinos LE, Coyne A, Schiantarelli J, Carlomagno Y, et al. Tau Protein Disrupts Nucleocytoplasmic Transport in Alzheimer’s Disease. Neuron. 2018;99(5):925–40.e7.

62. Cornelison GL, Levy SA, Jenson T, and Frost B. Tau-induced nuclear envelope invagination causes a toxic accumulation of mRNA in Drosophila. Aging cell. 2019;18(1):e12847.

63. Myeku N, Clelland CL, Emrani S, Kukushkin NV, Yu WH, Goldberg AL, et al. Tau-driven 26S proteasome impairment and cognitive dysfunction can be prevented early in disease by activating cAMP-PKA signaling. Nature medicine. 2016;22(1):46–53.

64. Zempel H, Luedtke J, Kumar Y, Biernat J, Dawson H, Mandelkow E, et al. Amyloid-β oligomers induce synaptic damage via Tau-dependent microtubule severing by TTLL6 and spastin. The EMBO journal. 2013;32(22):2920–37.

65. Zempel H, Thies E, Mandelkow E, and Mandelkow EM. Abeta oligomers cause localized Ca(2+) elevation, missorting of endogenous Tau into dendrites, Tau phosphorylation, and destruction of microtubules and spines. The Journal of neuroscience : the official journal of the Society for Neuroscience. 2010;30(36):11938–50.

66. Tian M, Chen Q, Graves AR, Goldschmidt HL, Johnson RC, Weinberger DR, et al. KIBRA-PKCγ signaling pathway modulates memory performance in mice and humans. 2021:2021.10.26.465926.

67. Crary JF, Shao CY, Mirra SS, Hernandez AI, and Sacktor TC. Atypical protein kinase C in neurodegenerative disease I: PKMzeta aggregates with limbic neurofibrillary tangles and AMPA receptors in Alzheimer disease. Journal of neuropathology and experimental neurology. 2006;65(4):319–26.

68. Amini N, Azad RR, Motamedi F, Mirzapour-Delavar H, Ghasemi S, Aliakbari S, et al. Overexpression of protein kinase Mζ in the hippocampus mitigates Alzheimer’s disease related cognitive deficit in rats. Brain Res Bull. 2021;166:64–72.

69. Höglund K, Schussler N, Kvartsberg H, Smailovic U, Brinkmalm G, Liman V, et al. Cerebrospinal fluid neurogranin in an inducible mouse model of neurodegeneration: A translatable marker of synaptic degeneration. Neurobiology of disease. 2020;134:104645.

70. VandeVrede L, Boxer AL, and Polydoro M. Targeting tau: Clinical trials and novel therapeutic approaches. Neuroscience letters. 2020;731:134919.

71. Chang CW, Shao E, and Mucke L. Tau: Enabler of diverse brain disorders and target of rapidly evolving therapeutic strategies. Science (New York, NY). 2021;371(6532).

72. Christensen DG, Meyer JG, Baumgartner JT, D’Souza AK, Nelson WC, Payne SH, et al. Identification of Novel Protein Lysine Acetyltransferases in Escherichia coli. mBio. 2018;9(5).

73. Collins BC, Hunter CL, Liu Y, Schilling B, Rosenberger G, Bader SL, et al. Multi laboratory assessment of reproducibility, qualitative and quantitative performance of SWATH-mass spectrometry. Nature communications. 2017;8(1):291.

74. Schilling B, Gibson BW, and Hunter CL. Generation of High-Quality SWATH(®) Acquisition Data for Label-free Quantitative Proteomics Studies Using TripleTOF(®) Mass Spectrometers. Methods in molecular biology (Clifton, NJ). 2017;1550:223–33.

75. Burger T. Gentle Introduction to the Statistical Foundations of False Discovery Rate in Quantitative Proteomics. J Proteome Res. 2018;17(1):12–22.

76. Mootha VK, Lindgren CM, Eriksson KF, Subramanian A, Sihag S, Lehar J, et al. PGC 1alpha-responsive genes involved in oxidative phosphorylation are coordinately downregulated in human diabetes. Nature genetics. 2003;34(3):267–73.

77. Subramanian A, Tamayo P, Mootha VK, Mukherjee S, Ebert BL, Gillette MA, et al. Gene set enrichment analysis: a knowledge-based approach for interpreting genome-wide expression profiles. Proceedings of the National Academy of Sciences of the United States of America. 2005;102(43):15545–50.

78. Shannon P, Markiel A, Ozier O, Baliga NS, Wang JT, Ramage D, et al. Cytoscape: a software environment for integrated models of biomolecular interaction networks. Genome research. 2003;13(11):2498–504.

79. Bindea G, Mlecnik B, Hackl H, Charoentong P, Tosolini M, Kirilovsky A, et al. ClueGO: a Cytoscape plug-in to decipher functionally grouped gene ontology and pathway annotation networks. *Bioinformatics (Oxford*, England). 2009;25(8):1091–3.

80. Moleirinho S, Chang N, Sims AH, Tilston-Lunel AM, Angus L, Steele A, et al. KIBRA exhibits MST-independent functional regulation of the Hippo signaling pathway in mammals. Oncogene. 2013;32(14):1821–30.

81. Ferguson L, Hu J, Cai D, Chen S, Dunn TW, Pearce K, et al. Isoform Specificity of PKMs during Long-Term Facilitation in Aplysia Is Mediated through Stabilization by KIBRA. The Journal of neuroscience : the official journal of the Society for Neuroscience. 2019;39(44):8632–44.

82. Montine TJ, Phelps CH, Beach TG, Bigio EH, Cairns NJ, Dickson DW, et al. National Institute on Aging-Alzheimer’s Association guidelines for the neuropathologic assessment of Alzheimer’s disease: a practical approach. Acta neuropathologica. 2012;123(1):1–11.

83. Mackenzie IR, Rademakers R, and Neumann M. TDP-43 and FUS in amyotrophic lateral sclerosis and frontotemporal dementia. Lancet Neurol. 2010;9(10):995–1007.

84. Tartaglia MC, Sidhu M, Laluz V, Racine C, Rabinovici GD, Creighton K, et al. Sporadic corticobasal syndrome due to FTLD-TDP. Acta neuropathologica. 2010;119(3):365–74.

85. Kim EJ, Sidhu M, Gaus SE, Huang EJ, Hof PR, Miller BL, et al. Selective frontoinsular von Economo neuron and fork cell loss in early behavioral variant frontotemporal dementia. Cerebral cortex (New York, NY : 1991). 2012;22(2):251–9.

86. McKhann GM, Knopman DS, Chertkow H, Hyman BT, Jack CR, Jr., Kawas CH, et al. The diagnosis of dementia due to Alzheimer’s disease: recommendations from the National Institute on Aging-Alzheimer’s Association workgroups on diagnostic guidelines for Alzheimer’s disease. Alzheimer’s & dementia : the journal of the Alzheimer’s Association. 2011;7(3):263–9.

87. Ljubenkov PA, Staffaroni AM, Rojas JC, Allen IE, Wang P, Heuer H, et al. Cerebrospinal fluid biomarkers predict frontotemporal dementia trajectory. Annals of clinical and translational neurology. 2018;5(10):1250–63.

